# Rho: A Purpose-Driven Workspace for Human–AI Collaborative Analysis

**DOI:** 10.64898/2026.11.03.763858

**Authors:** Yonghe Xia, Guangchuang Yu

## Abstract

Agent-assisted bioinformatics has moved the bottleneck from computing a result to judging it: whether a comparison answers the intended question, on which unit of replication, at which version of the input, and what would change the conclusion. These judgments need materials that are usually scattered across conversations, notebooks, figures, run logs and notes, while the agent’s context is a transcript rather than the structure the researcher is actually reading. We present Rho, a human–AI collaborative analysis environment that organizes materials by the current scientific purpose: it selects or obtains the materials a judgment needs, declares the relations they serve, composes operable views, and makes this organization a shared state that is at once the researcher’s reading interface and the agent’s continuation context. We present the design in three parts: a method object—the operations that form a round-trip, the kinds of state that must stay distinct, and the protocol invariants that make each distinction checkable; a three-layer architecture that separates capability providers from the application that composes them; and five illustrative analyses on public data. In a bulk RNA-seq case the recomputed counts differed from the published counts and the interface kept the computation, the choices and the discrepancy available for inspection; a single-nucleus case makes sample-level replication visible behind a cell-level comparison. Under one configuration these analyses show how the design behaves in practice; we present the work as a design contribution rather than an evaluation, and usability, comparative benefit and the scientific validity of individual analyses remain to be evaluated.

## Introduction

The bottleneck in agent-assisted bioinformatics is no longer computation but judgment. An agent can run a differential-expression comparison, but the researcher must still decide whether that comparison answers the intended question, on which unit of replication it rests, which version of the input produced it, and what would change the conclusion. A result can raise a new question, disagree with a published analysis, or turn on a choice— the comparison, the observation unit, the version of an input—and the next step depends on more than the latest answer. Researchers need to see the comparison, the data and code that produced it, the relevant results, and the choices that make an interpretation plausible or questionable. This paper argues that such judgments become tractable only when the materials and operations behind them are organized explicitly, made checkable, and shared by the person and the agent.

AI agents can write code and run analyses, but their outputs do not remove this need. An agent’s answer may summarize a result without exposing the observation unit, a changed input, or the analysis choices behind it. A conversation transcript can preserve the sequence of messages while leaving the researcher to reconstruct how a claim, file, calculation, and decision relate. Conversely, an analysis interface may make a figure visible without making the agent’s computational state or subsequent context accessible.

Two default arrangements do not answer this directly. One gives the interface to a workflow fixed at design time: views and controls are decided in advance and filled with content at run time. It is stable, but its organization is unrelated to a particular judgment— judging whether a difference is consistent across donors and judging why a computation did not finish show the same set of views. The other gives the process to a conversation: the agent writes, runs and explains, and the person reads the reply. It is flexible, but the material it forms is a byproduct of the dialogue; a person can read an explanation but cannot easily retrieve the material it rests on, and the agent’s next step depends mainly on the transcript rather than on the object the researcher is now examining.

Rho takes a third position: it makes the current scientific purpose the organizing principle, forms a working interface from it, and treats the organization itself as a fact shared by the person and the agent. The system obtains or selects the materials a judgment needs, states what each one contributes, composes the applicable views and operations, and points out what is still missing; the researcher reads, compares and revises within it, the agent continues from the same representation with its objects, scope and actual compute conditions, and new results return to the corresponding judgment. We use *discovering questions* broadly: a question may be supplied by the researcher, emerge from a discrepancy, or follow from an incomplete result. *Testing explanations* means making an explanation answerable to particular results and analysis choices; it does not mean that the interface can validate a biological mechanism by itself.

The proposed contribution is therefore a method, not a particular screen or a model capability: given the current scientific purpose and the capabilities actually connected, select or obtain materials from an ongoing analysis, organize their relations, and compose operable views, such that this organization is simultaneously the researcher’s interface for understanding the analysis and the agent’s shared context for continuing it, and is revised as human judgment and new results arrive. The claim is not that organization is a good idea but that its distinctions can be made checkable: the paper pairs each load-bearing claim with the protocol invariant that enforces it (Section 3.7). The paper makes three contributions.

1. **A method object**. We express “purpose ⍰ organization plan ⍰ operable interface ⍰ sourced shared state ⍰ agent continuation” as a set of checkable objects and operations, in which every step states its basis, gaps and version. Five operations form one round-trip; three kinds of change (layout, focus and analysis scope) must remain distinguishable; the shared state separates three sources (observed computation fact, agent interpretation, human judgment) and versions each object separately; each distinction is paired with the protocol invariant that enforces it (Table 2). A capability provider supplies information, compute and views; the organizer decides how they are composed.
2. **A system and five illustrative analyses**. Rho realizes this method in a three-layer architecture—application container, functional components and optional managed execution—connected by four shared concepts (action, resource, selection, view). We report five analyses on public data that illustrate different parts of the design under one configuration: a bulk RNA-seq discrepancy, a single-nucleus replication question, a spatial compartment comparison, an ageing-endothelium count discrepancy, and a patient-paired transcriptome comparison.
3. **A positioning**. We compare the method directly with the closest work, stating for each what it already provides, what this paper adds and what it does not claim, and identify the one question that those precedents leave open.

This is a design and method contribution: no experiment and no user study were run. Section 3 states the method; Section 4 describes the architecture; Section 5 reports the analyses; Section 7 sets the boundary and the criteria under which the claim fails. Figure 1 states the argument; Figure 2 the system that carries it; Figures 3 and 4 the evidence.

**Figure 1.**
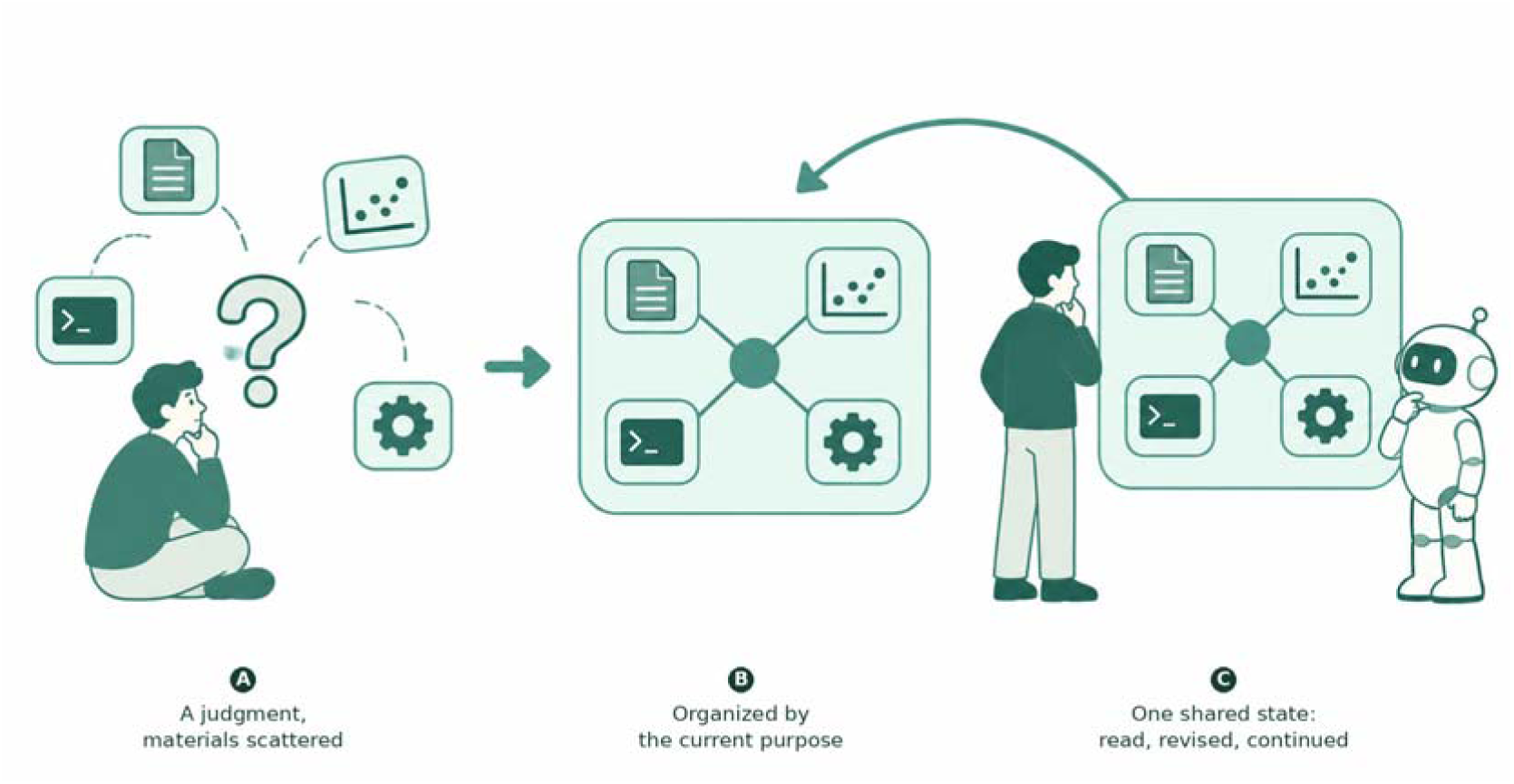
The argument. (A) A judgment whose materials are scattered across conversations, notebooks, figures and logs, while the agent’s context is a transcript rather than the structure the researcher is reading. (B) The materials are organized by the current purpose. (C) The organization is one shared state that the person reads and revises and the agent continues from. The protocol invariants that make these distinctions checkable are stated in Table 2.

## Related work

Research on interactive and agent-assisted analysis already provides several of the capabilities Rho brings together. We read each source at a recorded version and state what it establishes.

### Systems that make an analysis process inspectable

Making a generated analysis inspectable is an established line. WaitGPT organizes an agent’s code into a readable chain, shows execution state, and allows parameter changes, local follow-up questions and re-runs (Xie et al. 2024); interactive task decomposition exposes intermediate tasks for steering and verification and reports limits on how edits propagate (Kazemitabaar et al. 2024); AI Chains makes intermediate steps visible and editable (Wu, Terry, and Cai 2022); Treadstone renders the discussion of several analysis agents as a readable interface (Lee et al. 2026). InsightLens goes further and makes the insight itself the unit: it extracts and revises insights, links them to supporting code, outputs and figures, and navigates by theme (Weng et al. 2025). Loops uses provenance and visualization to support exploratory analysis in notebooks, including comparison across weeks (Eckelt et al. 2024); analytic provenance for sensemaking is a longer research agenda (Xu et al. 2015). These systems establish that a process can be read, that insights can be organized and revisited, and that inspection can pause. They organize by extracted conclusions or by the steps of a pipeline; they do not ask which materials a specific current judgment requires.

### Intent-driven interface generation

Intent-driven generation is a separate, older line. DynaVis synthesizes controls for visualization edits (Vaithilingam et al. 2024); DashChat composes dashboard prototypes (Shen et al. 2025); Generative UI produces whole pages from a prompt (Leviathan et al. 2026); Data Formulator 2 uses data threads to revisit states and supply context for later work (Wang et al. 2024). They rule out claiming intent-driven layout, or “a selected state as agent context”, as new. What they organize is a view, a dashboard or a page; the material a standing scientific judgment must bring together, and the relations among that material, are outside their scope.

### Shared scientific context, memory and revision

Sharing and revising scientific state also has precedents. Mycelium links scientific observations, interpretations and hypotheses into a sourced context graph and routes it to people and agents, and its supplementary figure records a human scientific constraint reaching a later analysis (Choudhury et al. 2026). SEMV localizes a challenge to a claim and its evidence, recomputes within the affected range, retains before-and-after reports, and consolidates verified experience into retrievable memory (Nguyen et al. 2026). Memory Sandbox offers memory that can be viewed, edited, summarized and traced to its source (Huang et al. 2023); cotomi Act turns a process into a workspace both sides can edit and an agent can retrieve (Oyamada et al. 2026); MemGPT brings external memory into the agent’s working loop (Packer et al. 2023). At the representation level we introduce no new provenance ontology: PROV-DM already expresses entities, activities, derivations and revisions (W3C 2013), and EVI expresses claims, evidence and challenges (Manir and Clark 2026). Thus “a challengeable explanation”, “retaining old results”, “local revision”, “revision entering memory” and “shared, editable memory” cannot be presented here as contributions.

### Statistical and bioinformatics constraints

The analysis setting rests on a scientific constraint that already has a literature: single-cell differential expression must account for biological replicates, and ignoring within-donor dependence produces false discoveries (Squair et al. 2021). Claim strength, the unit of replication and re-checking after a change have been raised directly for bioinformatics (Ang et al. 2026); interactive clarification of domain relationships and statistical models is the subject of Tisane and rTisane (Jun et al. 2022, 2024). This paper does not claim to discover pseudoreplication, to choose the right statistical model automatically, or that “claims need evidence” is a new idea. These works fix what a reader must understand; we are concerned with how such judgments arise, are retained, and change between reading and continuing an analysis.

### Plugin capability and heterogeneous compute

Connecting different compute and environments into one workbench also has mature foundations. Galaxy uses job-runner plugins and destination configuration to reach local, cluster and remote resources and sets environment conditions per destination (Galaxy Project 2026). JupyterLab’s extension points cover custom storage browsing, content and MIME rendering, run sessions and kernel services, and the application interface itself (JupyterLab 2026). Plugin architecture, customizable access, multiple compute backends and an extensible workbench therefore cannot on their own constitute this paper’s novelty.

### The remaining gap

Each line above solves one thing: the process can be read, insights can be organized, interfaces can be generated, state can be shared, capabilities can be plugged in. One question remains: **when all of these capabilities are present, what decides which materials should be organized into an operable working interface for the judgment at hand, and how does that organization become a fact shared by the person and the agent?** Existing systems decide the organization by a fixed workflow, by already-extracted insights, by a view intent, by a page, or by a transcript; none decides it by the materials and relations that a current scientific judgment requires. Table 1 places the closest work against this question and states the boundary of the claim explicitly.

**Table 1.**
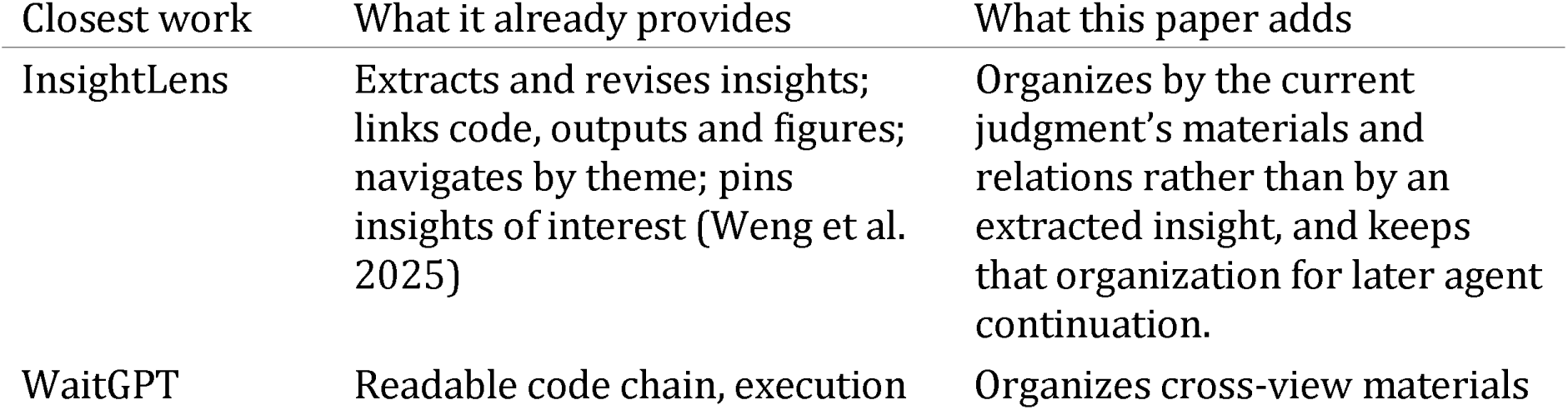
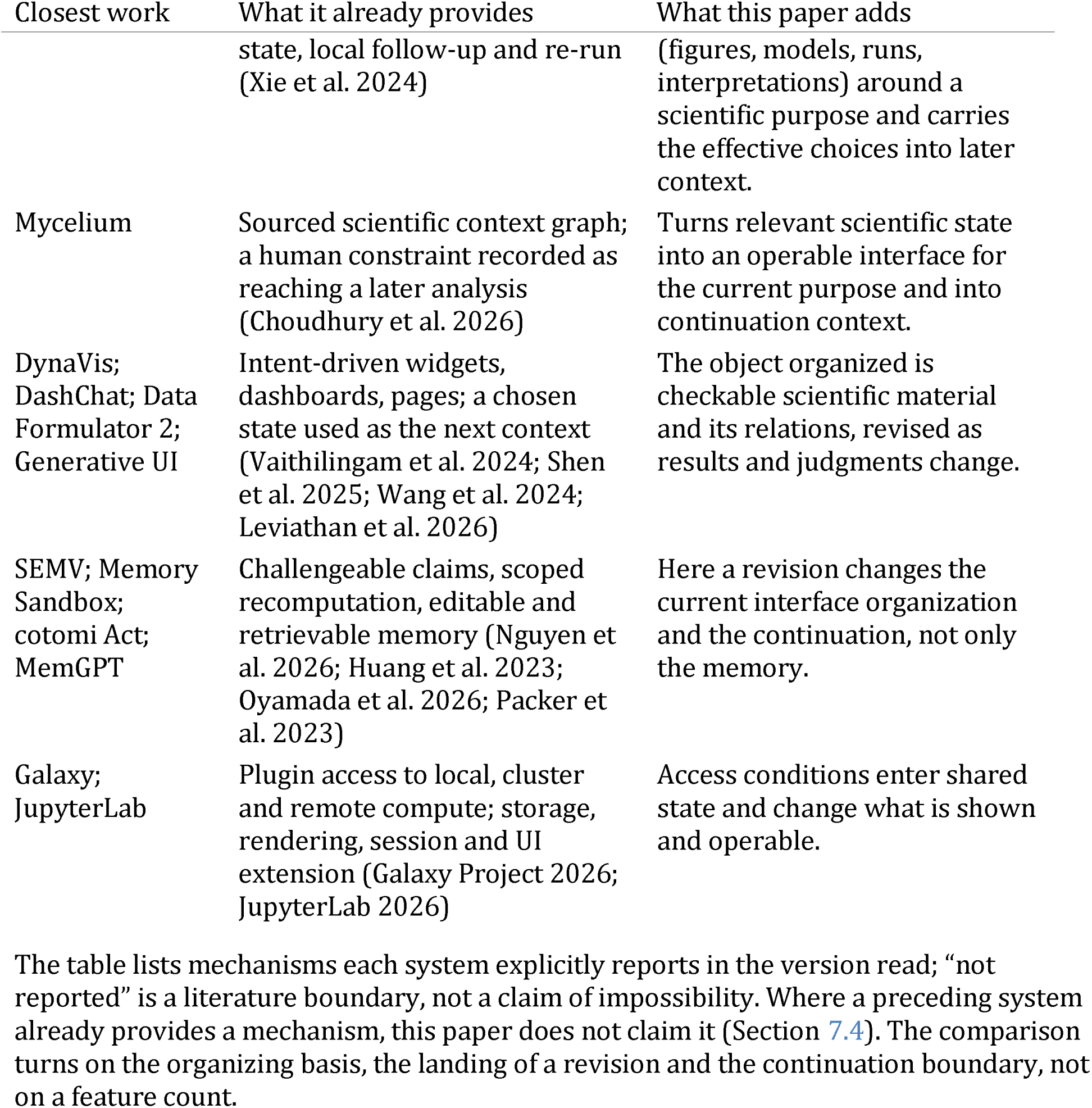
The closest work placed against the method’s question. Each cell reports a mechanism the source explicitly describes in the version read; the right column states what this paper adds.

| Closest work | What it already provides | What this paper adds |
|---|---|---|
| InsightLens | Extracts and revises insights; links code, outputs and figures; navigates by theme; pins insights of interest (Weng et al. 2025) | Organizes by the current judgment's materials and relations rather than by an extracted insight, and keeps that organization for later agent continuation. |
| WaitGPT | Readable code chain, execution | Organizes cross-view materials |
| Closest work | What it already provides | What this paper adds |
|  | state, local follow-up and re-run (Xie et al. 2024) | (figures, models, runs, interpretations) around a scientific purpose and carries the effective choices into later context. |
| Mycelium | Sourced scientific context graph; a human constraint recorded as reaching a later analysis (Choudhury et al. 2026) | Turns relevant scientific state into an operable interface for the current purpose and into continuation context. |
| DynaVis; DashChat; Data Formulator 2; Generative UI | Intent-driven widgets, dashboards, pages; a chosen state used as the next context (Vaithilingam et al. 2024; Shen et al. 2025; Wang et al. 2024; Leviathan et al. 2026) | The object organized is checkable scientific material and its relations, revised as results and judgments change. |
| SEMV; Memory Sandbox; cotomi Act; MemGPT | Challengeable claims, scoped recomputation, editable and retrievable memory (Nguyen et al. 2026; Huang et al. 2023; Oyamada et al. 2026; Packer et al. 2023) | Here a revision changes the current interface organization and the continuation, not only the memory. |
| Galaxy; JupyterLab | Plugin access to local, cluster and remote compute; storage, rendering, session and UI extension (Galaxy Project 2026; JupyterLab 2026) | Access conditions enter shared state and change what is shown and operable. |

The table lists mechanisms each system explicitly reports in the version read; "not reported" is a literature boundary, not a claim of impossibility. Where a preceding system already provides a mechanism, this paper does not claim it (Section 7.4). The comparison turns on the organizing basis, the landing of a revision and the continuation boundary, not on a feature count.

## A purpose-driven organization method

### Terms

- **Scientific purpose**. What the researcher is currently trying to judge or complete, for example “judge whether this difference is consistent across donors”. It may come from the current task and an explicit interaction context; a purpose inferred by the system should be visible and correctable.
- **Material.** An object that already exists or can be obtained in the analysis: data, annotations and mappings, figures, models, scripts, run records, outputs, literature evidence. Each material carries a version and a source.
- **Capability**. Connected information reading, computation and environment adaptation, with the corresponding views and operation entries. Capabilities are provided by plugins or native tools; their meaning belongs to the provider.
- **Analysis unit**. One analysis with its scientific question, scope, method and rationale, runs and artifacts, current interpretation, open items and revision history.
- **Organization plan**. The brief plan formed for a purpose this time: which objects and versions are selected, why they are needed, whether each serves comparison, explanation or further operation, what is still missing, and how the views locate the same object.
- **Shared state**. The versioned, attributed representation on which the person and the agent both rely; the interface and the agent’s context are different presentations of it.

In the implementation, an organization plan is carried by composition transactions and the shared state by the selection, annotation and prepared-context operations of the owners (Section 4); the two vocabularies name the same object at different levels.

### From a purpose to an organization plan

A purpose is first resolved into the relations that must be read or compared, and materials are chosen accordingly. Judging whether a difference is consistent across donors requires linking cells to donors, grouping to observations, descriptive material to statistical results; retrieving files that contain a gene name cannot do this organization. The system may propose these relations, but their basis must come from metadata, actual analysis records, or an explicit statement by either side. An organization plan handles four things: which materials are needed, where they can be obtained, what role each plays in the current judgment, and which views and operations are actually available. Before materials are juxtaposed, their observation units, inclusion scope and versions must be distinguishable; materials that cannot be compared directly can be used to explain a difference rather than forced into one result. If a required material does not exist, the plan states what is missing and which step remains, rather than filling in content to make the interface look complete; and it does not read “not selected” as “does not exist”.

### Three kinds of change must remain distinct

Operations that look similar on one interface have different meanings. Moving a view changes only layout; limiting the focus to one donor changes only the reading focus; changing the inclusion condition or the comparison changes the analysis scope. The system must not silently interpret the first two as the third, or a person’s reading action becomes an analysis decision and the agent receives a scope it never agreed to. Correspondingly, “collapsing code” is not “removing it from the analysis”, “closing the interface” is not “cancelling the analysis”, and “hiding in a view” is not “deleting from shared state”.

### Shared state: three sources and per-object versions

The content of the shared state is distinguished by source: the computation facts actually observed, the explanations and associations an agent gives, and the judgments a researcher explicitly writes. The three may reference one another but cannot substitute for one another: a successful exit is not a correct conclusion, an agent’s explanation is not a scientific fact, and a reading is not an adoption. Versions are per object: each analysis and each resource has its own version, with no global revision number. The person’s reading position, the analysis’s current version and the version the agent most recently received are kept separately; the same scope is not the same result, and receiving a revision is not recomputing or reading it.

### Five operations form a round-trip

One round-trip has five recognizable operations, not a mandatory pipeline. *Organize by purpose* selects or obtains the materials for the current judgment, states their roles and the gaps, and forms an operable layout. *Read and explore* expands, locates and compares them while keeping layout, focus and analysis-scope changes distinct. *Revise the purpose or the analysis* reselects materials or updates an interpretation and retains the original run facts and reading entry. *Continue from the shared state* supplies a checkable agent context and records the actual access state. Reorganize as work advances adds new results and states the change relative to what was read.

Revision happens only when the person explicitly changes an analysis choice or an interpretation state; asking “why” produces only an attributed explanation.

### Capability providers and the organizer

Plugins let the system adapt to different kinds of information, compute and environment without fixing one analysis stack at the base of the application. We distinguish the *capability provider* from the *organizer* of the working interface. A provider supplies a reading or compute entry, the actual format, scope, version and source it works with, and the views and operations that act on a declared object and scope; the organizer decides how they are composed. Three constraints follow. First, access conditions enter the shared state: which input is currently referenced, which capabilities are available, where computation runs, and which facts have not been obtained; otherwise “the interface looks complete” hides the actual gap. Second, a capability need not have a UI, and the catalog keeps five states apart—package presence, project enablement, dependency readiness, method authorization and UI renderability—so a missing view does not disable reading a table or editing a document. Third, a specific dataset, disease comparison, statistical design or analysis order is a project choice and must not be frozen into a general application merely because plugins are customizable. This division matches Rho’s source ownership (architecture, UI extension) and the plugin capability model; the five-state separation and the project-scoped selection are recorded in the headless collaboration record.

The method constrains only what must be checkable, not the layout, size or visual style. Its substantive constraints are: the selection and association of materials retain their attribution, and a model may propose but cannot turn a relation it generated into an established fact; the rationale and unresolved relations are traceable, and a person’s correction can target “a missing kind of material” rather than only a wrong conclusion; source identity, available operations, view bindings and existing state are carried by the corresponding capability rather than invented by the interface; and the state the interface rests on is the same set of versioned statements and references the agent receives, so compression cannot decide by a single free summary whether a human revision is kept.

### What makes the organization checkable

The method would be prose if nothing enforced its distinctions. Rho makes each load-bearing claim checkable through a specific protocol invariant, owned by a named mechanism and recorded in the implementation documents. The mechanisms the table names—composition transactions, selection channels, annotation anchors, prepared contexts—are defined in Section 4 and in the cited protocol records; here they appear only to show what each claim rests on. Table 2 states the claim, the invariant and the evidence this paper offers for it.

**Table 2.**
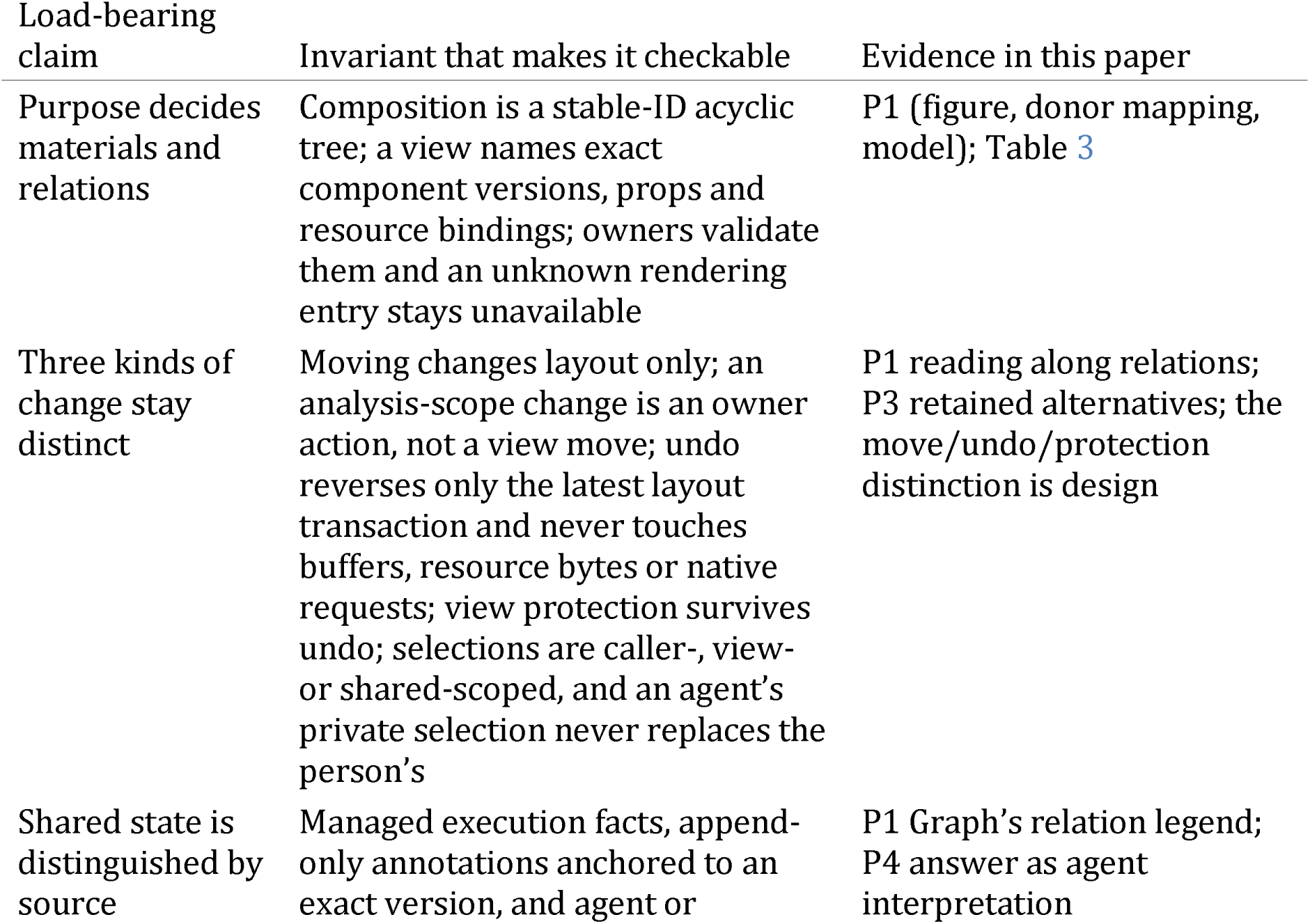
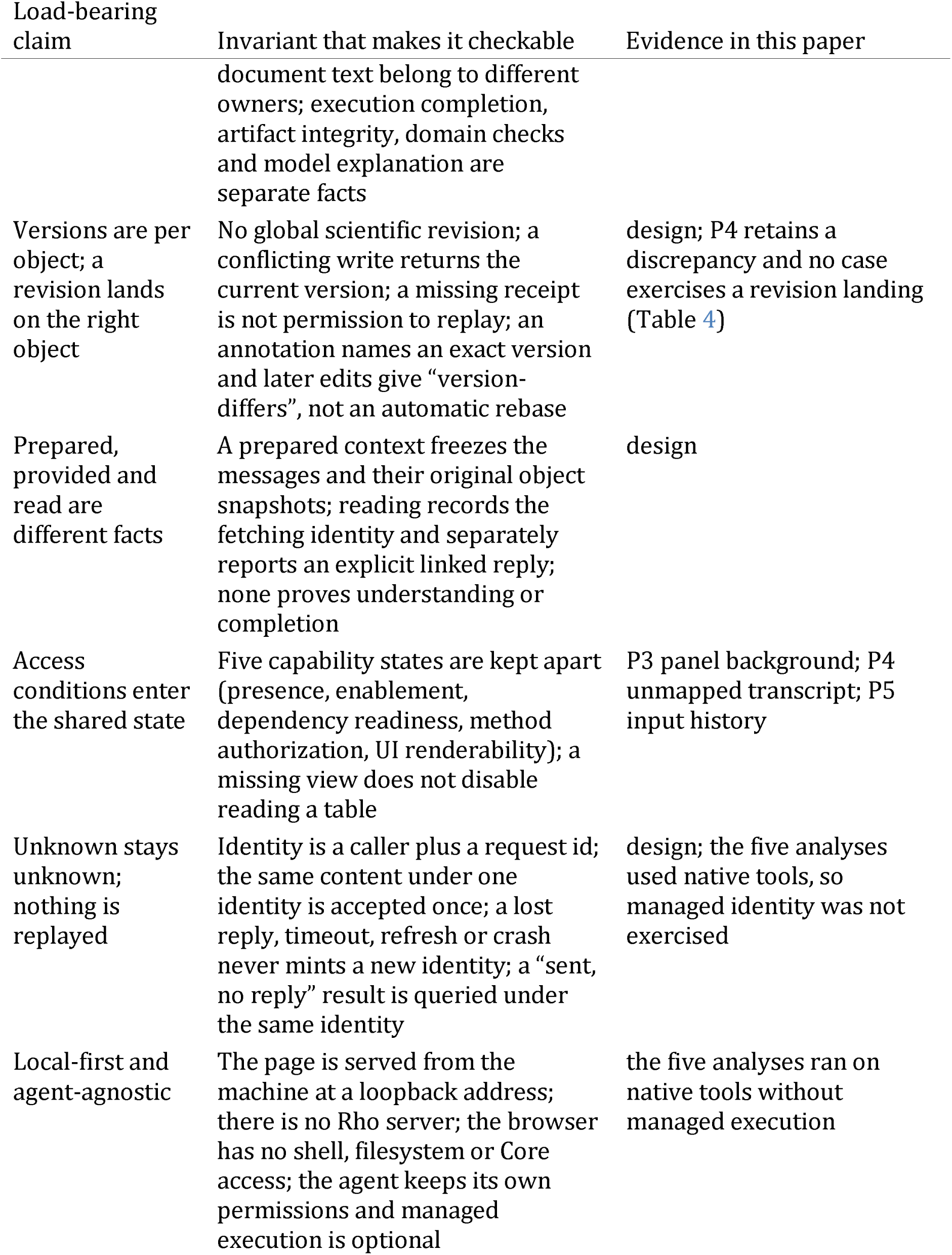
The argument and how it is checked. Each load-bearing claim is paired with the protocol invariant that makes it checkable and with the evidence offered here; “design” marks a claim the present analyses do not exercise.

| Load-bearing claim | Invariant that makes it checkable | Evidence in this paper |
|---|---|---|
| Purpose decides materials and relations | Composition is a stable-ID acyclic tree; a view names exact component versions, props and resource bindings; owners validate them and an unknown rendering entry stays unavailable | P1 (figure, donor mapping, model); Table 3 |
| Three kinds of change stay distinct | Moving changes layout only; an analysis-scope change is an owner action, not a view move; undo reverses only the latest layout transaction and never touches buffers, resource bytes or native requests; view protection survives undo; selections are caller-, view- or shared-scoped, and an agent’s private selection never replaces the person’s | P1 reading along relations; P3 retained alternatives; the move/undo/protection distinction is design |
| Shared state is distinguished by source | Managed execution facts, append-only annotations anchored to an exact version, and agent or document text belong to different owners; execution completion, artifact integrity, domain checks and model explanation are separate facts | P1 Graph’s relation legend; P4 answer as agent interpretation |
| Versions are per object; a revision lands on the right object | No global scientific revision; a conflicting write returns the current version; a missing receipt is not permission to replay; an annotation names an exact version and later edits give “version-differs”, not an automatic rebase | design; P4 retains a discrepancy and no case exercises a revision landing (Table 4) |
| Prepared, provided and read are different facts | A prepared context freezes the messages and their original object snapshots; reading records the fetching identity and separately reports an explicit linked reply; none proves understanding or completion | design |
| Access conditions enter the shared state | Five capability states are kept apart (presence, enablement, dependency readiness, method authorization, UI renderability); a missing view does not disable reading a table | P3 panel background; P4 unmapped transcript; P5 input history |
| Unknown stays unknown; nothing is replayed | Identity is a caller plus a request id; the same content under one identity is accepted once; a lost reply, timeout, refresh or crash never mints a new identity; a “sent, no reply” result is queried under the same identity | design; the five analyses used native tools, so managed identity was not exercised |
| Local-first and agent-agnostic | The page is served from the machine at a loopback address; there is no Rho server; the browser has no shell, filesystem or Core access; the agent keeps its own permissions and managed execution is optional | the five analyses ran on native tools without managed execution |

Protocol records: the headless collaboration and execution bridge documents define these invariants and operations, and Rho architecture defines ownership. The table maps claims to mechanisms and evidence.

## Rho: architecture and interface

### Three layers and four shared concepts

The method needs a carrier: Rho separates three layers (Figure 2). The **application container** selects and assembles components, and owns layout and shared interaction routing and presentation. **Functional components (plugins)** are the capability providers—an R analysis, an agent bridge, a table, a plot, an editor or a file tool—and own semantics, native state, artifacts, provenance and results. **Rho**-core supplies optional managed execution: original request identity and deduplication, retained execution facts, a resident Host and a public CLI. Core is not a prerequisite for an agent computation and is not on every path; an agent normally uses its native tools with existing permissions, and managed execution is selected only for a concrete need such as execution continuity or a retained record. During the five analyses reported in Section 5, managed execution was unavailable: they ran through the shared Notebook kernel on the agent’s native tools, so they exercise the workspace views and the shared session, not the optional managed-execution path. Rho is a local service with a browser interface: the page is served from the machine at a loopback address, there is no Rho server, and the browser has no shell, filesystem or Core access of its own—it calls named scoped operations on the local service. A project stays an ordinary folder, an agent client keeps its own permissions, and the scientific materials stay on the machine.

Four concepts connect components to the container: an **action** (identity, typed inputs, preconditions, effects, results and cancellation semantics); a **resource** (an owner-provided reference to a file, table, plot, model or task, with provenance and the content version it needs); a **selection** (stable entities or ranges in a particular resource version); and a **view** (presentation of resources, registered actions and observation subscriptions). Agent tools, UI controls and CLI adapters reach the same component action, so a person and an agent reach the same capabilities and resources through different entrances. In the design, the shared state the method relies on is this body of actions, resources, selections and views, each with an owner, a version, and a provenance whose source is an observed computation fact, an agent interpretation, or a human judgment (Figure 1C). Two facts the figure keeps apart: component semantics belong to the plugin, composition and routing to the application, and a single execution’s lifetime to Core; and a displayed relationship is not by itself proof that a scientific claim is correct. The five analyses below instantiate views and resources over a shared Notebook session; they do not exercise the full action and selection contract described here.

**Figure 2.**
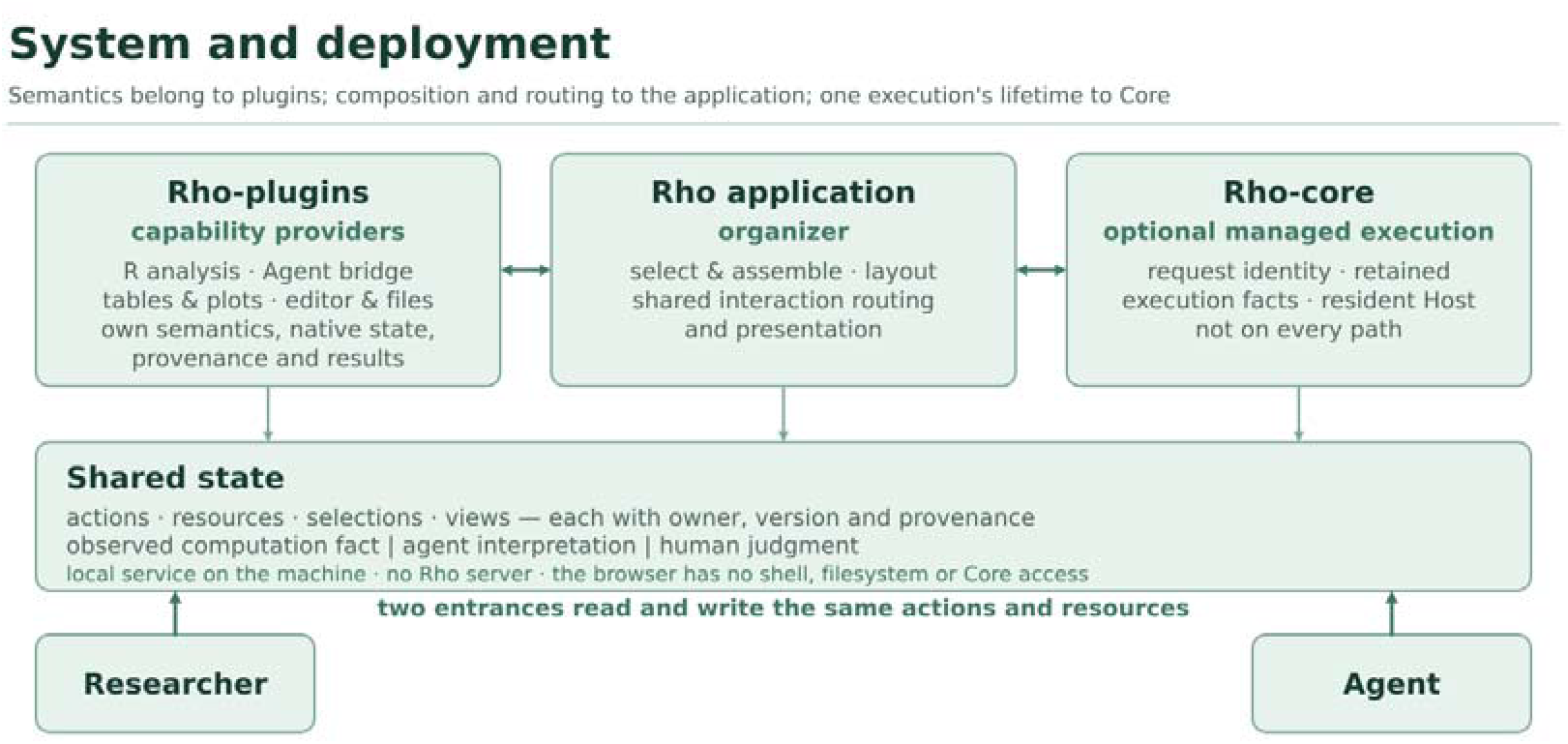
System and deployment. Semantics stay with capability providers (R analysis, the agent bridge, tables, plots, editors), composition and routing with the application, and one execution’s lifetime with an optional core; the researcher and the agent reach the same actions and resources. The service is local: there is no Rho server, and the browser has no shell, filesystem or Core access.

These boundaries have a practical consequence for the method. Because the application composes views and routes actions but does not re-implement a component’s behavior, the same analysis objects can appear in different views without a second source of truth. Because Core is optional, an analysis can run with the agent’s native tools while still being observed and represented in the shared state. And because a missing component is reported unavailable rather than silently installed, the interface can state a gap instead of hiding it.

### Execution identity and unknown outcomes

For work that asks for execution continuity, Rho reaches Rho-core only through its public CLI, and the application side of that boundary mints a request identity once—a caller name plus a request id—and keeps it with the action. Submitting the same content again returns the original run and dispatches nothing; different content under the same identity is a conflict; a lost reply, a wait timeout, a browser refresh, a view failure or a service crash never mints a new identity. A “sent but no reply” result is treated as possibly executed: the system queries the same identity and shows the actual facts, including an unknown outcome, instead of starting fresh work. A resident host is held independently of tabs, views and agent sessions, so closing a view releases observation and rendering only. This is the concrete form of the paper’s claim that prepared, provided and read are different facts and that an unobserved outcome stays unknown; the exit-code mapping that makes “do not replay” mechanical is recorded in the execution bridge document.

### The workspace views

Rho organizes work around a project containing analysis materials such as data, scripts, notes, outputs and recorded activity. The **Notebook** provides an R session in which the agent’s cells, outputs and live variables can be inspected. **Studio** brings selected materials and conversation views together; the **Overview** presents an answer with its associated files; **Graph** displays associations drawn from notes and activity. These are views of particular recorded states, not a guarantee that all materials are semantically linked or that an agent has read every displayed item.

The guiding design question is how a researcher can move from a question to the evidence needed to assess an explanation, and back to the next analysis. The required materials differ by question: checking a reversal count calls for contrasts and inclusion rules, whereas assessing a single-nucleus comparison calls for sample membership as well as a cell embedding. Rho also distinguishes material of different origins: an agent’s summary or stated plan, a note, a command reference, an observed file and a computational output support different inferences. Reading a displayed result is not equivalent to endorsing it, and a further analysis may change a result without retroactively making an earlier interpretation valid. The interface provides entry points for inspection; scientific judgment and appropriate statistical design remain the researcher’s responsibility.

### Case selection and evidence

We describe five selected analyses using publicly available GEO datasets associated with published biomedical studies. They were run on one local machine with one agent and one model in Rho 0.5.2 (source commit 64d73550f4bb1786e6cec42071fbe8a67a8ba230), with managed execution unavailable; the public data were downloaded before the runs and R code was executed in the shared Notebook session. The five are drawn from a corpus of 97 fully open papers. The implementation record reports that two of the five needed one turn and three needed two to four, because the agent client capped at 64 turns or waited on an event that nothing wakes, with 8–24 approval choices per run and resource paths guessed. These are operational facts of this configuration, cited to the status and literature-reproduction records, not properties of the method. The tasks cover particular expression analyses rather than complete reproductions of the studies. The local project scripts, result tables, notes, session exports and screenshots underpin the descriptions, but their implementation record is not yet publicly available. The original studies and GEO accessions identify the input sources; they are not independent evidence that these local analyses are correct.

We report the first case in depth because it exercises all three workspace views; the other four are reported more briefly as different questions that call for different evidence. Table 3 summarizes the five. All are observational demonstrations of what the interface displayed and what the saved analyses reported; there was no independent participant or comparative user study. We distinguish screenshots captured during the original P1 session from later *3840×1972* captures of reopened P2–P5 projects, where the saved views and artifacts were available but the original R sessions had ended. Separately redrawn plots are identified in their captions.

**Table 3.** The five illustrative projects and the question used to examine each one. Each task covers only a selected part of its source study.

| Case | Public data | Analysis | Question for inspection |
|---|---|---|---|
| P1 | GSE274459 | Bulk RNA-seq | How do comparison and reversal definitions relate to the differing counts? |
| P2 | GSE221215 | Single-nucleus RNA-seq | How do sample replication and cell annotation affect interpretation? |
| P3 | GSE232853 | GeoMx spatial profiling | Which compartment contributes to a pathway interpretation? |
| P4 | GSE270697 | Bulk RNA-seq | Which input and analysis choices could relate to the reported discrepancy? |
| P5 | GSE252990 | Blood transcriptome | What can a patient-paired comparison support? |

Where Table 2 reads claim-first, Table 4 reads evidence-first: for each case, which part of the method it illustrates and what it leaves untested.

**Table 4.** What each case illustrates of the method, and what it does not show. The mapping is uneven: no single case exercises the full round-trip or a revision landing.

| Case | Part of the method it illustrates | What it does not show |
|---|---|---|
| P1 | Organize and read across the three workspace views; keep the three material origins distinct (agent cell, note, observed output) | A human revision or an agent continuation; any user observation |
| P2 | Reading that separates the cell count from the biological replicate; an agent annotation kept as interpretation, not fact | A completed round-trip or a revision landing; the class labels are unvalidated |
| P3 | Retaining alternative analysis choices as inspectable material; reading a pooled enrichment for its missing direction | A revision or continuation; no comparison with the source study |
| P4 | Keeping a count discrepancy and its candidate causes open, with no tested cause | Which choice caused the difference; the metabolomics arm |
| P5 | Distinguishing a paired comparison from a different question; sensitivity to input handling | Any revision or agent continuation; the original processing history |

### Illustrative analyses

Across the five cases the round-trip is uneven, and Table 4 marks which part of the method each case illustrates. We report P1 in depth, because it exercises the workspace views and shows one judgment made inspectable (Figure 3); the other four are reported more briefly as a set of judgments that turn on different material relations (Figure 4). None of the five closes a human revision into an agent continuation; that part of the design is described in Section 4 and is demonstrated, as a separate episode, in Section 6.

### P1. A discrepancy in tauopathy bulk RNA-seq

The first project started from a published bulk RNA-seq study of chromogranin A and tauopathy and its GEO data (Jati et al. 2025). The requested task was narrower than reproducing the paper: recompute specified differential-expression and reversal counts, perform GO biological-process enrichment, and record outputs and choices. The public data had been downloaded before the run. A local project README stated the comparisons and expected files; the 17 samples comprised five wild-type, six hTau and six CgA-KO/hTau samples. The agent executed R code in Rho’s shared Notebook session, using two pairwise DESeq2 contrasts with nominal P < 0.01 and log2 fold change > 0.5.

The first result was a discrepancy, not a confirmation. The local run reported 1,096 up- and 1,549 down-regulated genes for the first contrast, versus 1,193 and 869 reported in the task’s comparison with the paper. The reversal counts were 334 and 551, versus 417 and 293. The recorded analysis retained alternative reversal definitions in a separate table. These counts describe one run and one comparison; they do not establish why the published numbers differed, nor do they establish which definition is scientifically correct. Figure 3B sets the two sets of counts side by side.

**Figure 3.**
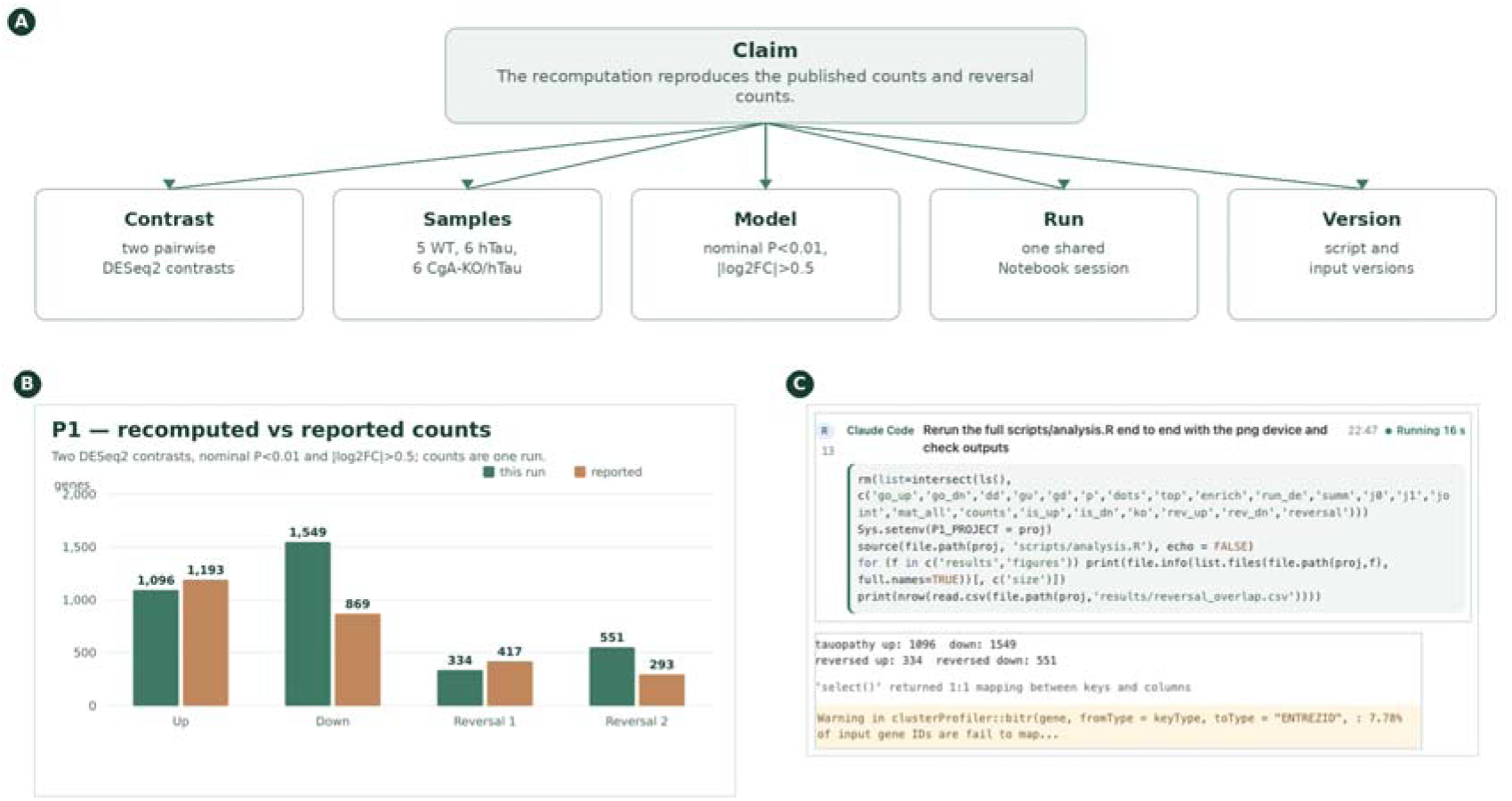
One P1 judgment, made inspectable. (A) The claim and the materials it rests on: the contrast, the samples, the model, the run and the version. (B) The recomputed and reported counts, which make inspection necessary. (C) The real Notebook session: an agent-attributed cell and its reported counts. The judgment is inspectable because these materials and versions are explicit, not because its counts are right.

The discrepancy changes what the researcher needs to inspect. Rather than asking only whether the run produced a figure, the researcher can ask which samples and contrast were used, how reversal was defined, which choices affected the counts, and whether those choices match the published analysis. This is the method’s first operation made concrete: the purpose is not “produce a figure” but “explain a difference”, and the organization must keep the computation and its materials available for that investigation.

#### Inspecting the computation

Figure 3C shows two regions of the P1 Notebook capture: the agent-attributed R cell and its reported output.

The Notebook made it possible to inspect an agent-attributed cell alongside its output. The same session also held data frames, plotting objects and enrichment results (Figure S1). This is an observed property of the run, not evidence that the analysis was correct or that the person understood it.

#### Bringing the analysis back together

Beyond the Notebook, two further views carry the same project. Studio places the written question next to the materials produced during the work; Graph shows recorded associations among a question, its findings, files and runs (Figures S2 and S3).

Studio identifies the data and method, lists files, and preserves the unresolved difference in counts, so a conclusion can be traced to a script, output or recorded choice without treating a generated summary as the only source of truth. Graph distinguishes several kinds of evidence: an agent’s report that a result exists, a filename mentioned in a command, an observed disk change and a statement in notes. Keeping them distinct helps a reader see what still requires checking.

**Figure 4.**
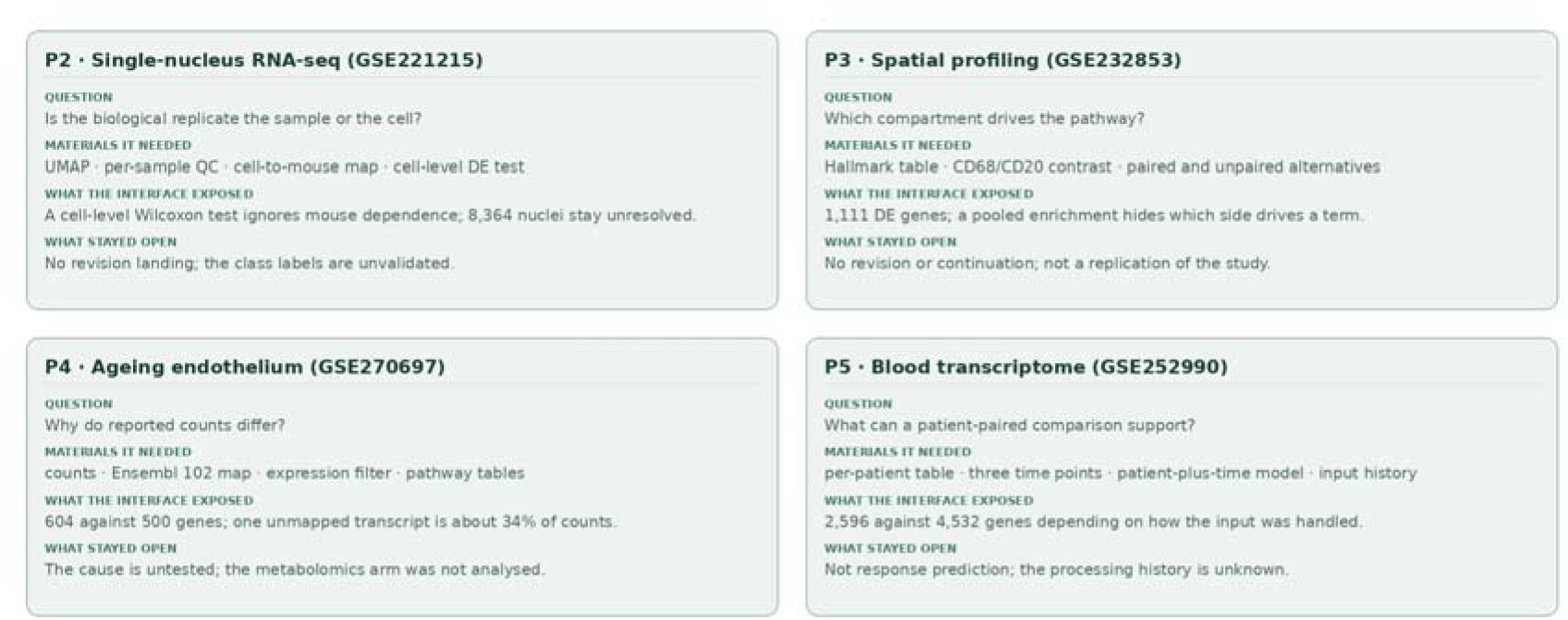
Four further judgments. P2–P5, each turning on a different material relation: the biological replicate, the compartment behind a pathway, the cause of a count difference, and what a paired comparison can support. The cards state what the interface exposed and what stayed open; none closes the round-trip.

### P2. Single-nucleus data: is the biological replicate the sample?

The second project used hippocampal single-nucleus data from a study of neuronal APOE4 removal (Koutsodendris et al. 2023) (GEO GSE221215). The local task began with 11 samples and retained 92,597 nuclei after quality control; the agent clustered the nuclei and assigned coarse cell classes, 8,364 of which remained unresolved. These labels are working annotations, not independently validated cell identities. The saved embedding and per-sample quality-control table are in Figure S4; Figure 4 (top left) states what the case exposed.

The visual grouping opens a specific question. The local excitatory-neuron comparison contained 20,832 PS19-fE4 and 8,533 PS19-fE3 nuclei from four and three mice, respectively, and its cell-level Wilcoxon test did not model dependence among nuclei from the same mouse; a large number of nuclei is not the same replication as a large number of mice (Squair et al. 2021). The per-sample table exposes sample contributions while the UMAP shows how the assigned classes occupy the embedding; neither establishes that the differential-expression list is reliable. The comparison also asks a different question from the study’s central neuronal APOE4-deletion intervention, so the gene list was not compared with the paper’s. A defensible continuation would validate the class labels, align the intended contrast with the study, and repeat the comparison with an appropriate sample-aware model. Rho makes the materials for that decision accessible; it does not supply the biological judgment.

### P3. Spatial profiling: which compartment supports an explanation?

The third project used the GeoMx digital spatial profiling dataset GSE232853 from a diffuse large B-cell lymphoma study (Liu et al. 2024). The local matrix contained 16,560 genes and 578 areas of illumination, including CD68- and CD20-selected segments. The task compared those two compartments, first checking marker expression and then fitting a limma model with `duplicateCorrelation` for paired segments, and it retained paired-only and unpaired alternatives so that the choice of comparison stayed inspectable. The significant Hallmark terms are re-plotted in Figure S9.

At adjusted P < 0.05 and log2 fold change > 0.58, the main analysis reported 1,111 differentially expressed genes, of which 833 were higher in CD68 and 278 higher in CD20. An over-representation analysis of the combined list identified 13 significant Hallmark sets, including complement and interferon-gamma response. Because the analysis pooled both directions, an enriched term does not say which compartment drives it; the relevant next question is directional. The recorded panel background also matters: these results describe enrichment within the measured gene panel, not an unrestricted genome-wide universe. The local matrix did not include the spatial-region labels required for the original study’s region-level comparison, so this is not a replication of the full spatial study.

### P4. Ageing endothelium: why do reported counts differ?

The fourth project examined the transcriptomic arm of an ageing study that also included metabolomics (Kiesworo et al. 2025) (GSE270697). The local analysis used three old and four young mouse skeletal-muscle endothelial samples; after transcript-to-gene mapping and filtering it fitted a limma–voom comparison. At BH-adjusted P < 0.05 and log2 fold change > 1 it reported 604 genes (370 higher and 234 lower in old samples), whereas the project records the paper’s corresponding total as 500 (406 higher and 94 lower). These figures make the discrepancy visible without explaining it (Figure S10).

The saved choices identify several questions a reviewer can pursue: the local analysis used an Ensembl 102 transcript map and a particular expression filter where the paper used a different pipeline, and one transcript absent from the candidate annotations accounted for approximately 34% of counts in each sample, with its effect on normalization unmeasured. These identify potential sources of difference, not a tested cause. The pathway view shows why the underlying table matters: KEGG oxidative phosphorylation had a negative normalized enrichment score ( 2.74) for old versus young, and all terms in the displayed selection are lower in old samples, which should not be read as the direction of every significant pathway. The metabolomics arm was not analysed in this project.

### P5. Patient transcriptomes: a paired comparison, not response prediction

The fifth project used the blood-expression component of a multi-omics mesothelioma study (Chin et al. 2025) (GSE252990). Its local matrix contained measurements from 40 patients at three time points; at baseline, 23 were recorded as responders and 17 as non-responders, whereas the study’s phase II cohort was reported as 31 and 23. These are different scopes: the available matrix does not represent the entire 54-person cohort. Figure 4 (bottom right) summarizes the case; the saved result table is Figure S7.

For the on-treatment question, the analysis paired baseline and C3D1 samples from the same 40 patients using a patient-plus-time design. The available matrix contained continuous, apparently normalized values, so the main local analysis applied log2(x + 1) and limma `lmFit`/`eBayes` and reported 2,596 genes at BH-adjusted P < 0.05, while an alternative that applied voom directly to the supplied values reported 4,532. The contrast between outputs exposes sensitivity to input handling; it is not evidence that treating normalized values as counts is appropriate, or that either count is a verified biological finding. The pairing addresses within-patient change and asks a different question from baseline prediction of response, and it does not by itself distinguish treatment effects from other changes over time. The useful continuation is to establish the matrix’s processing history, check the model and sample metadata, and compare the appropriate contrast with the original study.

### Collaboration beyond the Rho window

This section carries the continuation part of the argument: how work that a person and an external agent share becomes visible in the project and continues from one state. An agent need not originate in Rho for a researcher to bring its work into the project. In an installed external ACP episode (Rho contributors 2026b), an existing Codex session used an explicitly configured relay: the original client kept its input, permission choices and runtime, while Rho displayed the session’s reported plan and references to project files. Rho can also observe an external Claude Code or dsh session read-only through a project-scoped collector: it shows tool requests and their returns, refusals, native exit codes and output files, keeps them while the view is open, and marks a request with no observed return as awaiting a result. Reconnection reads the same stream after its existing cursor; it does not start a host, restart a collector, submit work or resume a model, and opening the view sends no message and starts no agent or host. Outputs are observed as files with current paths, byte counts and SHA-256 prefixes; they are not automatically attributed to a tool call or declared complete, and a captured snapshot is labelled “producer unconfirmed” until an owner says otherwise. Opening a snapshot inserts an exact resource and version into the workspace through the same conditional-composition path as any other view: a stale layout is refused and protected views are not replaced. A filename or a current hash does not by itself establish authorship or scientific correctness. This is a specific relay-based connection, not passive attachment to an arbitrary running client (Figure S8); the observation boundary is recorded in the agent-activity document.

A separate base-R collaboration episode (Rho contributors 2026a) demonstrates the scientific handoff itself. An automated operator selected May in an `airquality` table and asked an external Claude Code agent why there were 24 observations and to exclude only missing ozone values. The agent received the selected row and version through a project-scoped MCP handoff, changed the script and linked the recalculated outputs. May changed from 24 observations with mean ozone 24.125 to 26 with mean 23.615; an independent base-R expression checked the five monthly aggregates, and the original and new tables remained separately available for comparison. This is a controlled demonstration with an automated operator, not evidence of human usability or a validation of the missing-data assumption. The project’s own review of that run is more specific: the screens did not present the answer, the change between the two results had to be found by comparing two tables, and implementation facts stood where the question and its answer should be. That is an unresolved presentation defect of this configuration rather than a property of the method, and it is recorded in the product-direction record.

Rho’s bundled PA14 quorum-sensing project (Rho contributors 2026c) provides another starting point. It brings together swarming measurements, co-culture ratios, RNA-seq counts and annotations to ask what the las and rhl systems contribute to collective behaviour. It is an illustrative project, not a sixth independent analysis in the five-project episode.

## Discussion

Rho’s design makes an analysis available as a connected body of work: a question, its materials, computational activity, results and recorded choices. The five cases give the method a concrete use: a mismatch with published counts, the biological replicate behind a cell-level comparison, direction in a spatial contrast, sensitivity to annotation, and a patient-paired comparison each demand different evidence. The external-agent episodes show two routes by which work initiated elsewhere can be connected to project materials.

### Difference from the closest work

The comparison in Table 1 is meant to be read as three different landing points rather than one slogan. Relative to InsightLens, the organizing unit moves from an extracted insight to the materials and operations a current judgment requires. Relative to WaitGPT, the working interface links a scientific judgment to the computation that is meant to answer it, not only to the code chain. Relative to Mycelium, relevant scientific state is further turned into an interface a person can read and operate while an agent continues from the same state. The difference cannot be shortened to “InsightLens only displays, WaitGPT only shows code, Mycelium only has a back end”.

Mycelium is especially close, and could even host this method’s storage and routing. Rho should therefore not claim to invent shared scientific memory; the difference is how a working interface is formed for a purpose, and how that interface’s organization and interaction take part in a continuing analysis. If an implementation merely adds a summary and a browsing page on top of such a graph, the difference does not hold.

### When the claim does not hold

Several criteria can directly defeat the candidate contribution:

- If the same collaboration can be obtained by adding “automatic notes a generated dashboard an appended summary to the model” to an existing system, then prior work already covers it and there is no independent increment.
- If “the current purpose” can only degrade into a search keyword or a fixed template, and material selection and relation statements do not produce an organization a person can check, then Section 3 has no substance.
- If the interface organization cannot distinguish layout change, focus change and analysis-scope change, then a person’s reading action is mistaken for an analysis decision and the shared state cannot be trusted.
- If a revision lands only by timestamp on the latest version rather than by object identity and declared relations, then a revision during parallel progress falls on the wrong scope.
- If the access conditions (local, remote, results-only) do not enter the shared state, the interface presents the incomplete as complete.

These criteria also point to the work needed: implement the round-trip on a real project, observe whether a person therefore judges evidence more easily and whether the agent therefore continues on the right object, and check whether the cost of organizing errors, layout jitter and relation maintenance is controlled.

### Costs and risks

The method carries concrete costs: the system may misread a purpose, select the wrong material or hide a counterexample; an organization that changes with purpose may interrupt reading; extracting and scoping scientific relations adds burden; a single entrance may obscure differences between data and environments; an organization that is easy for a person to read is not necessarily the best context for an agent; and the runtime may not expose an observable point of attachment. Selection, conditions, gaps and updates must therefore stay checkable, and automatic generation cannot be used to make these problems disappear. “A reasonable layout” is a design goal that has to be argued for; it does not follow from automatic generation or from the availability of features.

### Limitations and future evaluation

The examples use one configuration and selected, pre-existing datasets. They do not compare Rho with an alternative interface, measure users’ understanding or decision quality, or establish full reproduction of the source studies; no benchmark of analysis outcomes is reported. The P2 cell classes and comparisons require sample-aware validation; P3’s combined enrichment has no direction; P4’s unexplained annotation loss and count discrepancy need further investigation; and P5 depends on input-processing choices. No single case shows that Rho improves a scientist’s judgment, and independent researcher use has not been observed; the status record makes no usability claim for the same reason. The checks in Section 3.7 are protocol-level: they show that a distinction is enforced, not that it improves a judgment.

The next evidence should include an accessible, versioned record of each analysis; independent scientific review of the scripts, inputs and interpretations; and observations of researchers pursuing their own questions. A comparative study could test whether the interface helps people locate the evidence behind a claim, identify a discrepancy or choose a suitable next step.

## Conclusion

When computation and agents are no longer the bottleneck, human judgment still needs materials, relations and operations to be organized. This paper presents a design that makes that organizing work explicit: select materials by the current scientific purpose, state their relations, compose operable views, bring the actual access conditions into the same representation, and let this organization be both the researcher’s reading interface and the agent’s shared continuation context, with human judgment and new results moving along the same set of versioned objects. The single-cell and bulk analyses show where the mechanism touches a scientist’s work concretely, and where it meets the precedents of generative interfaces, shared scientific context, memory revision and plugin-based access. The paper does not claim any of those capabilities individually; it argues for the link that remains unexplained when they are combined, and that link stands or falls on whether it can be observed on a real project, not on a list of features.

## Data, code and evidence availability

The public source datasets are GEO GSE274459, GSE221215, GSE232853, GSE270697 and GSE252990. The Rho implementation and the published external-agent episode records are available at https://github.com/YuLab-SMU/Rho; the analyses used Rho 0.5.2 at source commit 64d73550f4bb1786e6cec42071fbe8a67a8ba230, and the bundled PA14 project is cited separately. The five analysis projects and their execution record are local and are not yet publicly archived. Their versioned release, including scripts and outputs sufficient to assess the stated results, is necessary before this manuscript can support independent review of those claims.

## Notes

### Competing Interest Statement

The authors have declared no competing interest.

### Funding

- National Natural Science Foundation of China: 32670859

