## Supplementary Figures for "Rho: A Purpose-Driven Workspace for Human–AI Collaborative Analysis"

Yonghe Xia Guangchuang Yu

This supplement holds the full interface captures behind the manuscript's five analyses. Each is a screenshot of one recorded state under one configuration (Rho 0.5.2, one agent, one model, one local machine, managed execution unavailable). A capture shows what the interface displayed; it does not by itself establish that an analysis was correct, that a displayed relationship is scientifically valid, or that a person understood or endorsed an output. Regions are rearranged or cropped for legibility; pixel content is unchanged.

### Supplementary Figures

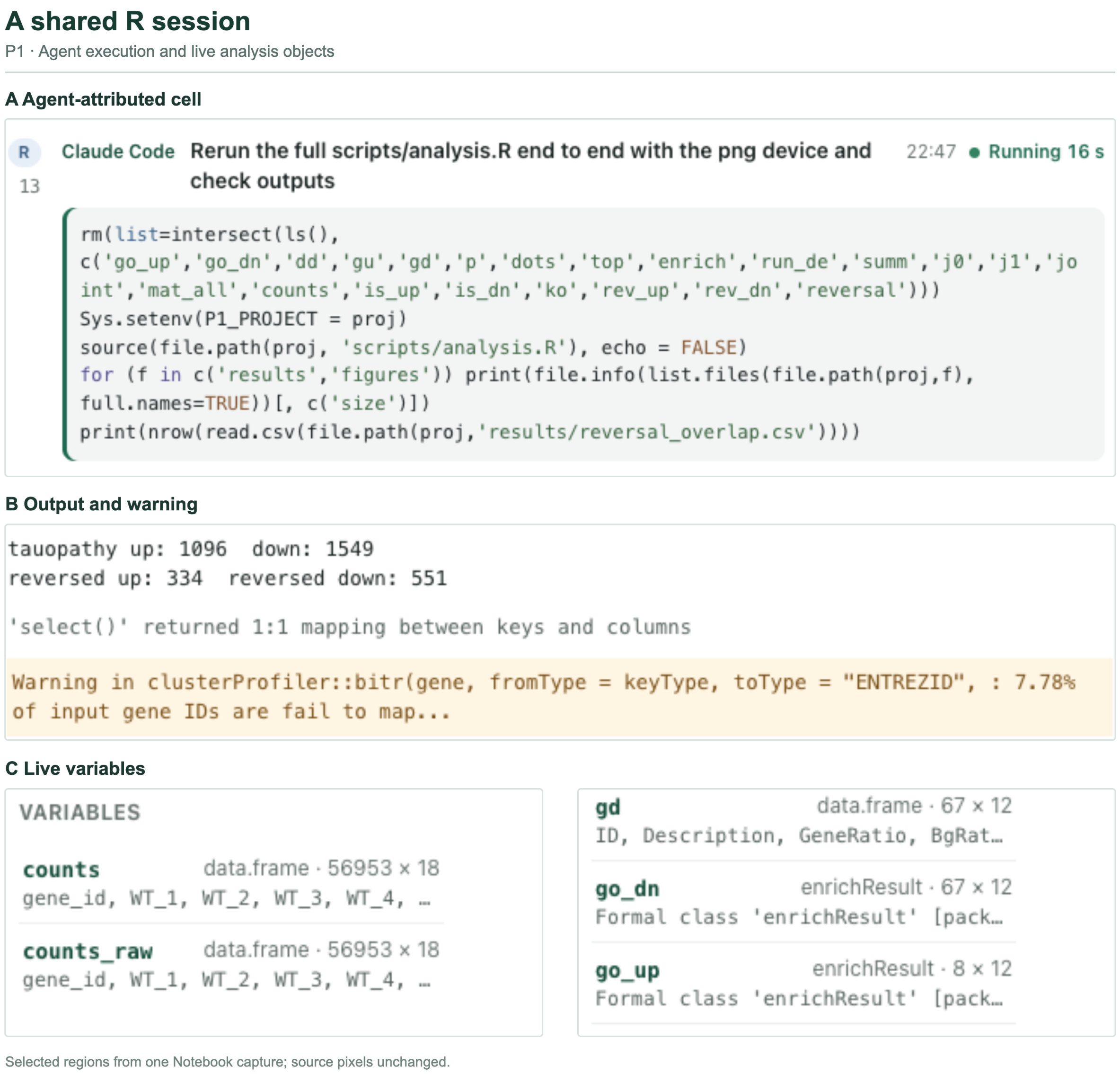


Figure S1: **The P1 Notebook (full capture).** (A) An R cell attributed to Claude Code, with its stated purpose and elapsed time; (B) reported counts and a gene-mapping warning; (C) two parts of the live Variables pane, including `enrichResult` objects. Regions rearranged for legibility.

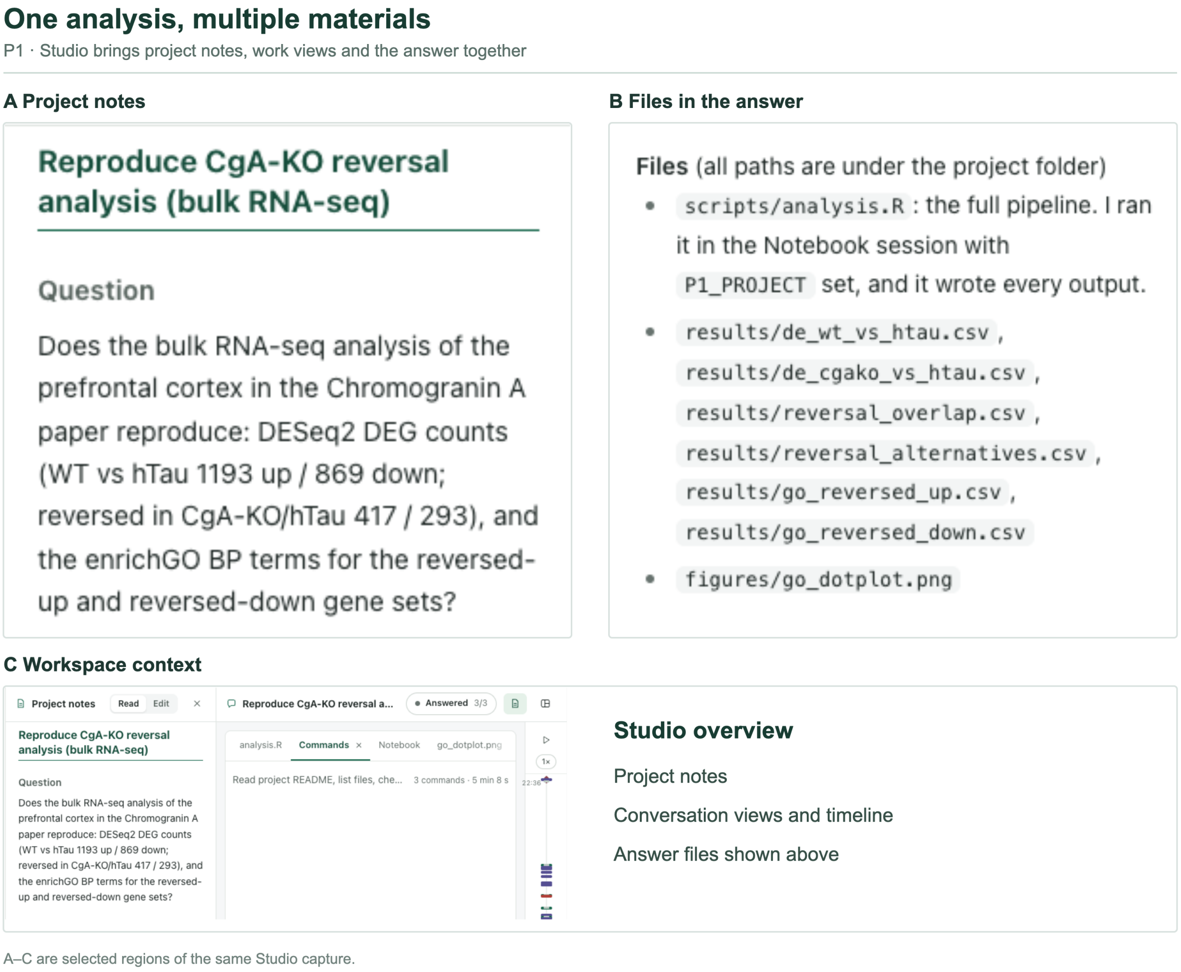


Figure S2: **P1 in Studio.** (A) The question in project notes; (B) files listed in the agent's answer; (C) the conversation's work views and timeline. Presence of a component does not establish that the agent read it.

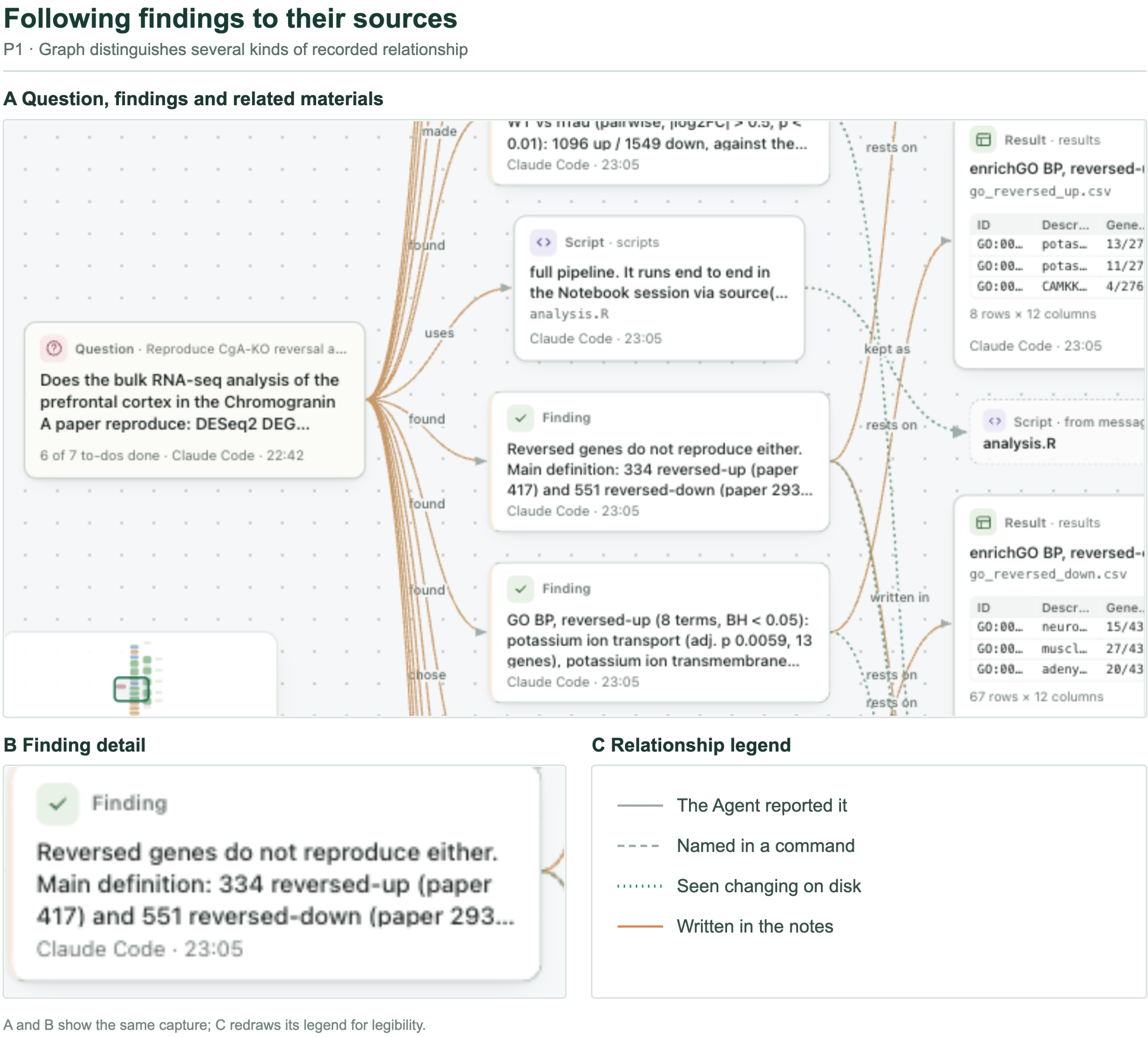


Figure S3: **P1 in Graph.** (A) A graph region relating a question, findings, script and result files; (B) an enlarged finding detail; (C) the relationship legend, redrawn for legibility. Edges make recorded associations navigable; they do not certify a scientific explanation.

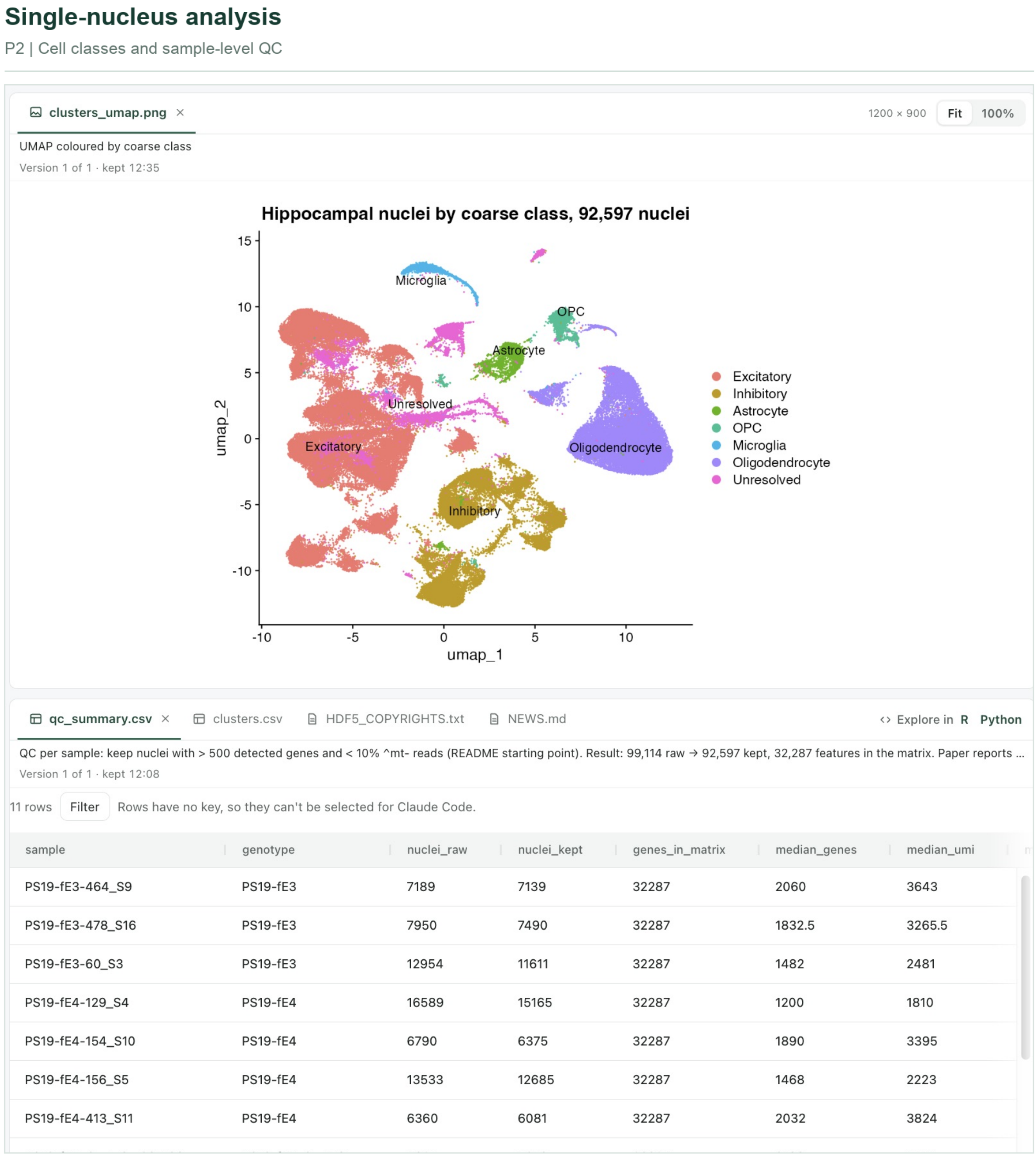


| sample | genotype | nuclei_raw | nuclei_kept | genes_in_matrix | median_genes | median_umi |
|---|---|---|---|---|---|---|
| PS19-fE3-464_S9 | PS19-fE3 | 7189 | 7139 | 32287 | 2060 | 3643 |
| PS19-fE3-478_S16 | PS19-fE3 | 7950 | 7490 | 32287 | 1832.5 | 3265.5 |
| PS19-fE3-60_S3 | PS19-fE3 | 12954 | 11611 | 32287 | 1482 | 2481 |
| PS19-fE4-129_S4 | PS19-fE4 | 16589 | 15165 | 32287 | 1200 | 1810 |
| PS19-fE4-154_S10 | PS19-fE4 | 6790 | 6375 | 32287 | 1890 | 3395 |
| PS19-fE4-156_S5 | PS19-fE4 | 13533 | 12685 | 32287 | 1468 | 2223 |
| PS19-fE4-413_S11 | PS19-fE4 | 6360 | 6081 | 32287 | 2032 | 3824 |

Figure S4: **P2 in Studio.** The saved UMAP and per-sample QC table, reopened after the run (source capture $3840 \times 1972$). The classes are preliminary and the R session was no longer live at capture.

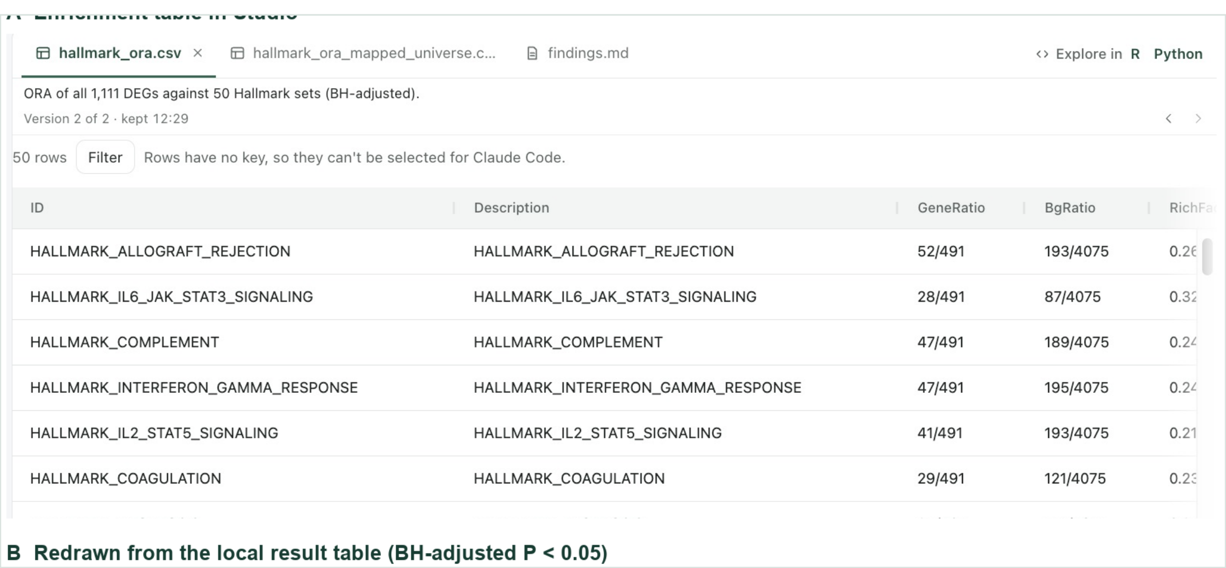


Figure S5: **P3 in Studio.** The saved Hallmark over-representation table as it appeared in the reopened project (3840 × 1972 capture). The 13 significant terms are re-plotted in Figure S9; the redraw is not a new enrichment analysis.

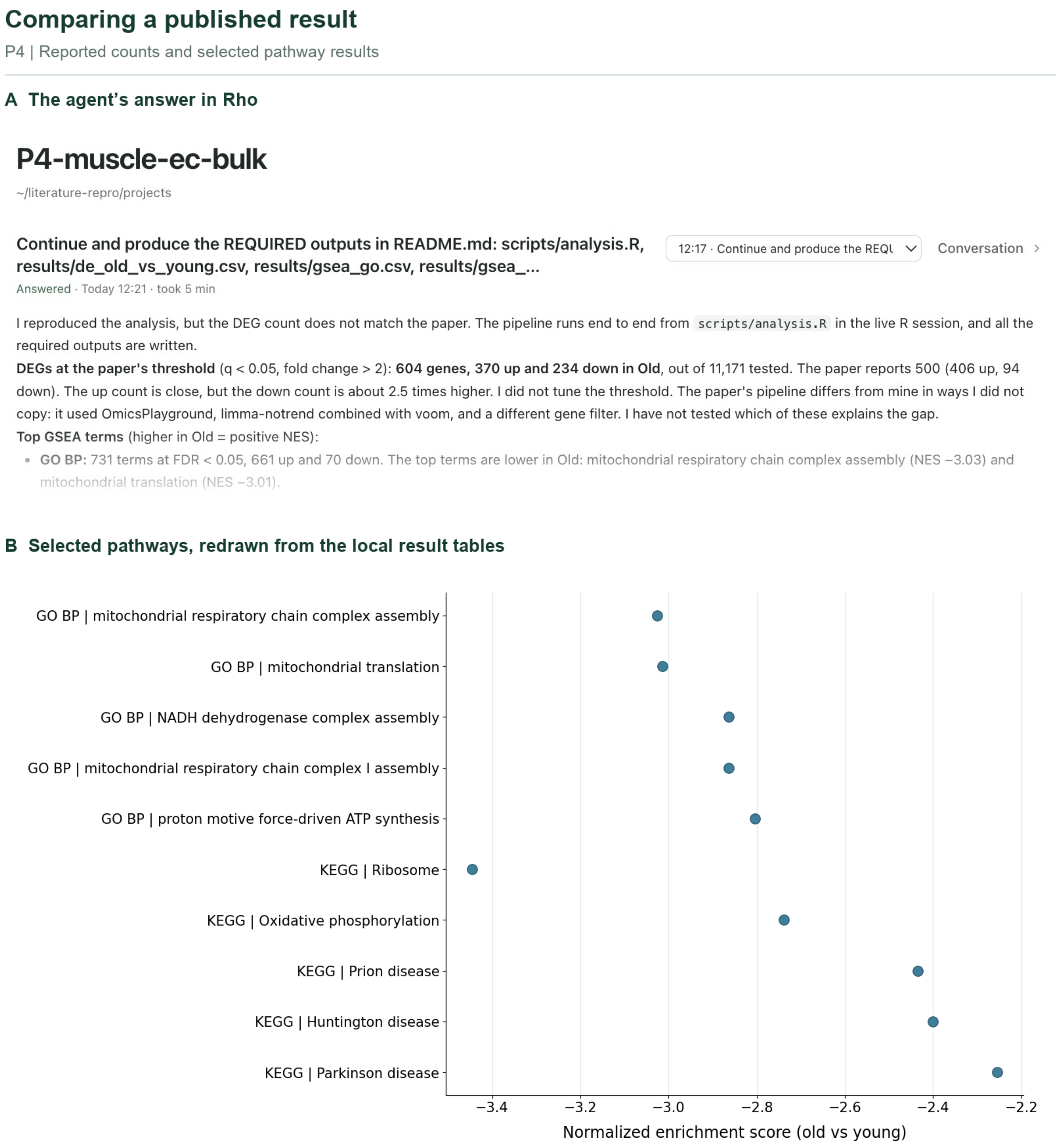


Figure S6: **P4 overview and selected pathways.** (A) The agent's reported counts; (B) a redraw of the five lowest-adjusted-P terms from each local GO BP and KEGG table. The answer is the agent's interpretation, not independent validation.

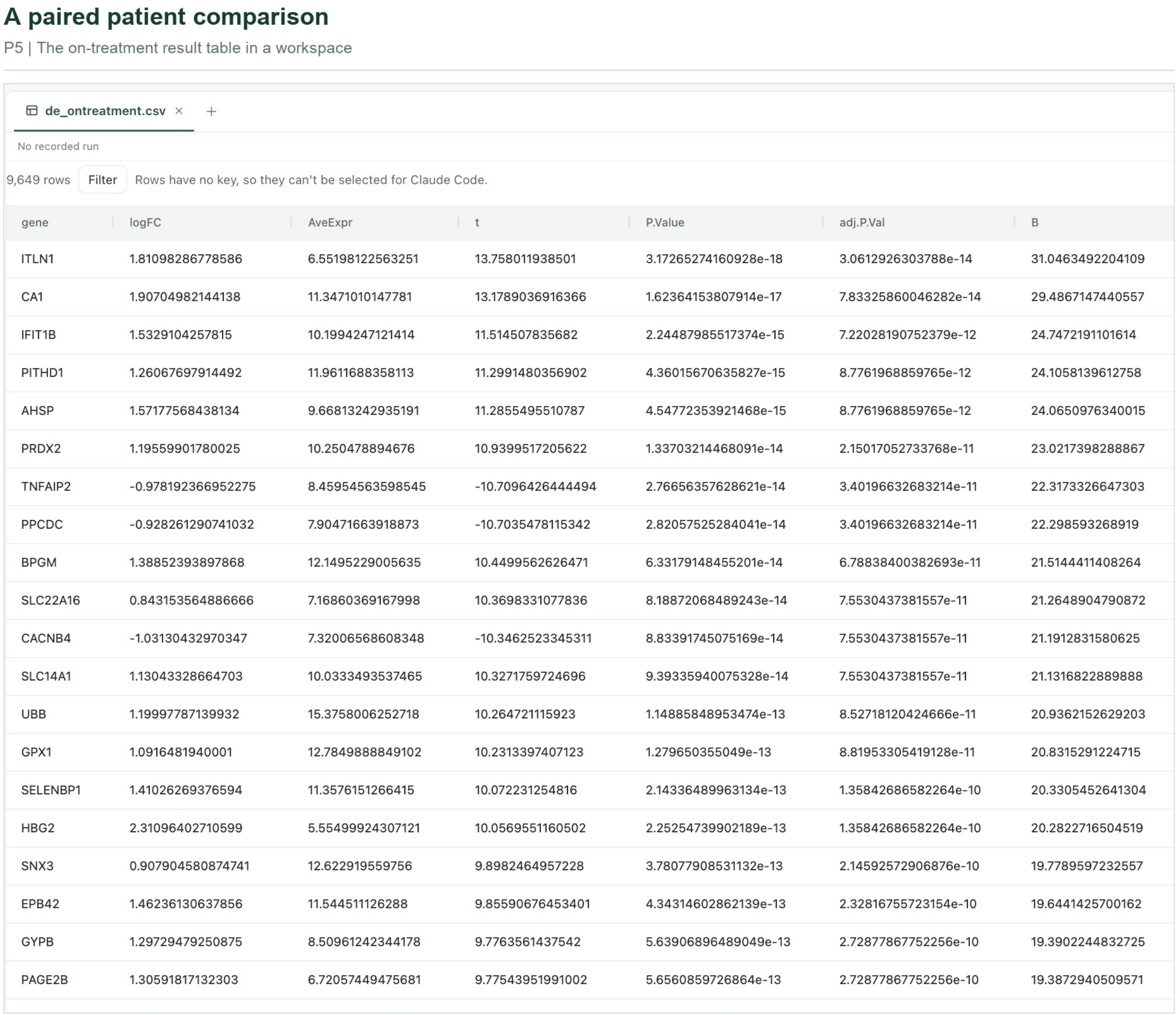



| gene | logFC | AveExpr | t | P.Value | adj.P.Val | B |
|---|---|---|---|---|---|---|
| ITLN1 | 1.81098286778586 | 6.55198122563251 | 13.758011938501 | 3.17265274160928e-18 | 3.0612926303788e-14 | 31.0463492204109 |
| CA1 | 1.90704982144138 | 11.3471010147781 | 13.1789036916366 | 1.62364153807914e-17 | 7.83325860046282e-14 | 29.4867147440557 |
| IFIT1B | 1.5329104257815 | 10.1994247121414 | 11.514507835682 | 2.24487985517374e-15 | 7.22028190752379e-12 | 24.7472191101614 |
| PITHD1 | 1.26067697914492 | 11.9611688358113 | 11.2991480356902 | 4.36015670635827e-15 | 8.7761968859765e-12 | 24.1058139612758 |
| AHSP | 1.57177568438134 | 9.66813242935191 | 11.2855495510787 | 4.54772353921468e-15 | 8.7761968859765e-12 | 24.0650976340015 |
| PRDX2 | 1.19559901780025 | 10.250478894676 | 10.9399517205622 | 1.33703214468091e-14 | 2.15017052733768e-11 | 23.0217398288867 |
| TNFAIP2 | -0.978192366952275 | 8.45954563598545 | -10.7096426444494 | 2.76656357628621e-14 | 3.40196632683214e-11 | 22.3173326647303 |
| PPCDC | -0.928261290741032 | 7.90471663918873 | -10.7035478115342 | 2.82057525284041e-14 | 3.40196632683214e-11 | 22.298593268919 |
| BPGM | 1.38852393897868 | 12.1495229005635 | 10.4499562626471 | 6.33179148455201e-14 | 6.78838400382693e-11 | 21.5144411408264 |
| SLC22A16 | 0.843153564886666 | 7.16860369167998 | 10.3698331077836 | 8.18872068489243e-14 | 7.5530437381557e-11 | 21.2648904790872 |
| CACNB4 | -1.03130432970347 | 7.32006568608348 | -10.3462523345311 | 8.83391745075169e-14 | 7.5530437381557e-11 | 21.1912831580625 |
| SLC14A1 | 1.13043328664703 | 10.0333493537465 | 10.3271759724696 | 9.39335940075328e-14 | 7.5530437381557e-11 | 21.1316822889888 |
| UBB | 1.19997787139932 | 15.3758006252718 | 10.264721115923 | 1.14885848953474e-13 | 8.52718120424666e-11 | 20.9362152629203 |
| GPX1 | 1.0916481940001 | 12.7849888849102 | 10.2313397407123 | 1.279650355049e-13 | 8.81953305419128e-11 | 20.8315291224715 |
| SELENBP1 | 1.41026269376594 | 11.3576151266415 | 10.072231254816 | 2.14336489963134e-13 | 1.35842686582264e-10 | 20.3305452641304 |
| HBG2 | 2.31096402710599 | 5.55499924307121 | 10.0569551160502 | 2.25254739902189e-13 | 1.35842686582264e-10 | 20.2822716504519 |
| SNX3 | 0.907904580874741 | 12.622919559756 | 9.8982464957228 | 3.78077908531132e-13 | 2.14592572906876e-10 | 19.7789597232557 |
| EPB42 | 1.46236130637856 | 11.544511126288 | 9.85590676453401 | 4.34314602862139e-13 | 2.32816755723154e-10 | 19.6441425700162 |
| GYPB | 1.29729479250875 | 8.50961242344178 | 9.7763561437542 | 5.63906896489049e-13 | 2.72877867752256e-10 | 19.3902244832725 |
| PAGE2B | 1.30591817132303 | 6.72057449475681 | 9.77543951991002 | 5.6560859726864e-13 | 2.72877867752256e-10 | 19.3872940509571 |

Figure S7: **P5 in Studio.** The saved on-treatment versus baseline patient-level table, reopened after the run; the historical notebook and project records are needed to inspect its analysis context.

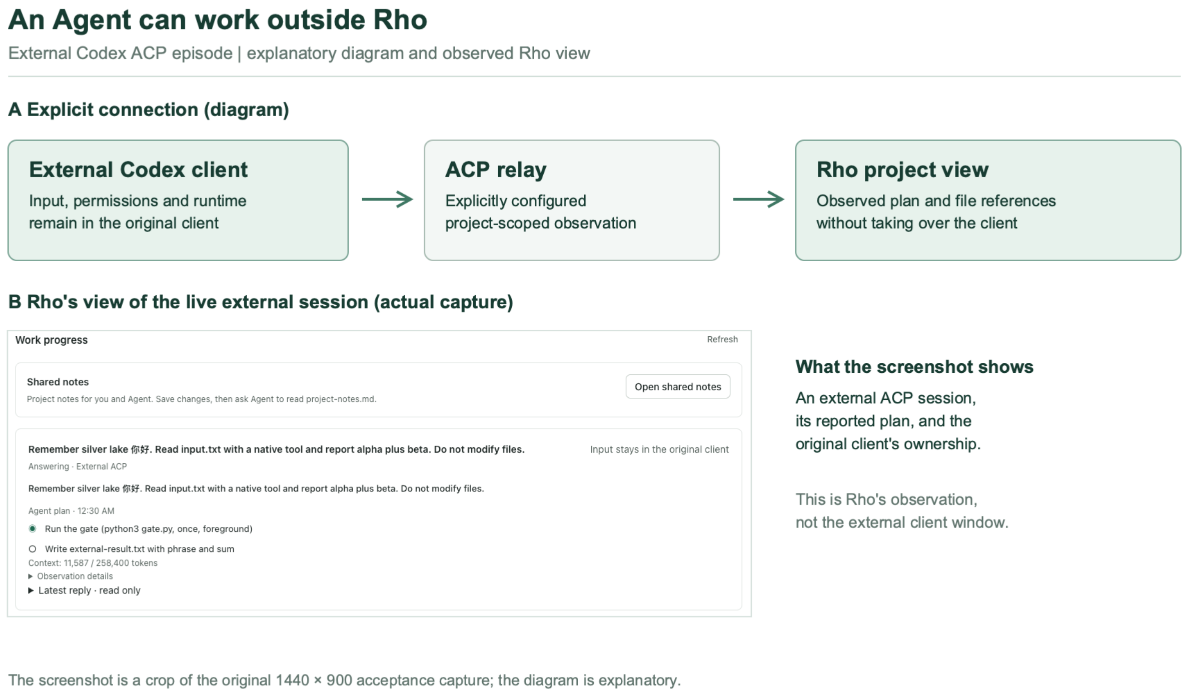


Figure S8: **An external agent and Rho.** (A) An explanatory diagram of the explicit ACP connection; (B) a Rho-side capture during the separate installed external Codex episode, showing a live reported plan (original $1440 \times 900$); it does not show the external client window. This episode used a different package composition from the five 0.5.2 analysis projects.

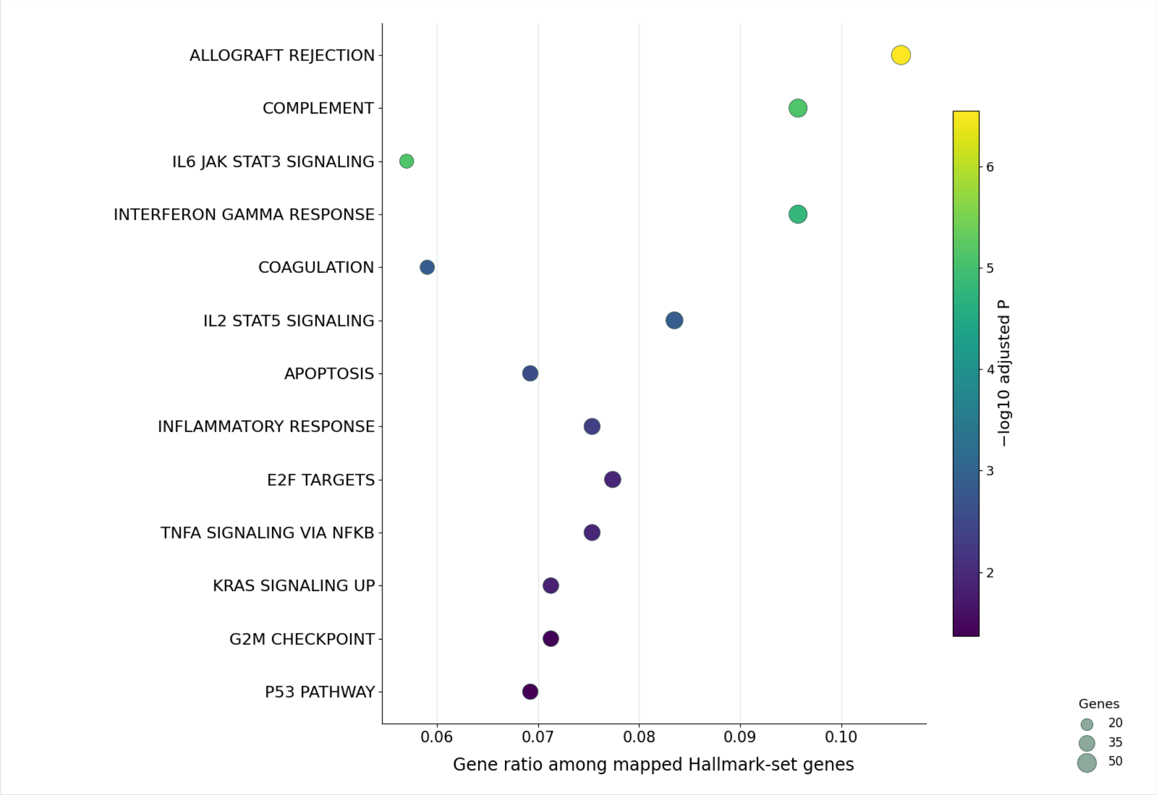


Figure S9: **P3 Hallmark terms.** A redraw of the local Hallmark over-representation table: the 13 terms with Benjamini–Hochberg-adjusted P $< 0.05$. Point position is the recorded gene ratio, size the number of genes, colour the adjusted P. The redraw uses the same local table and is not a new enrichment analysis; the source table as it appeared in Studio is Figure S5.

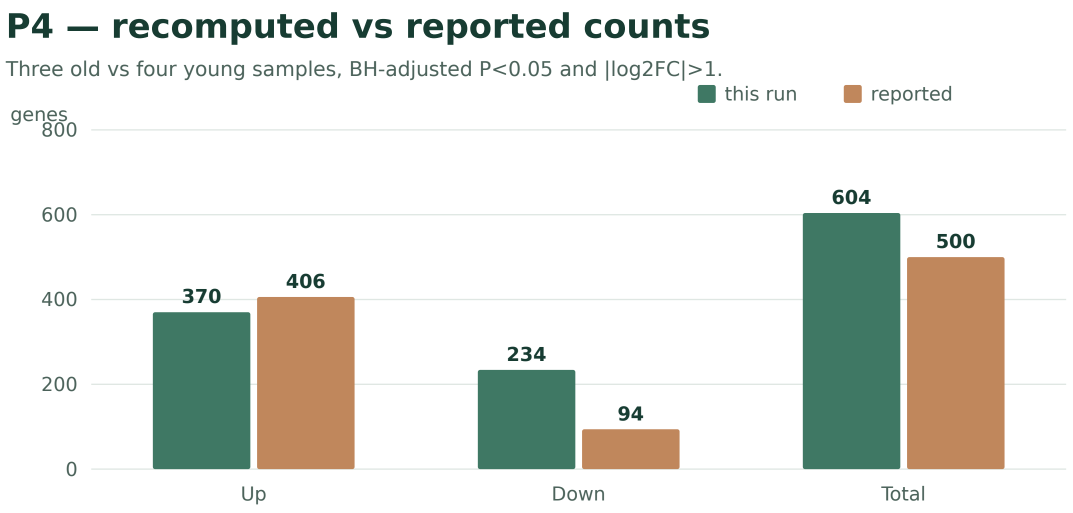


Figure S10: **P4 recomputed versus reported counts.** The local run reports 604 genes (370 higher, 234 lower in old samples) against the paper's recorded 500 (406 higher, 94 lower). The chart redraws the counts given in the text; it is not a new analysis and does not identify the cause. The agent's overview and its selected pathways are in Figure S6.
